# Plasma-Enabled Multiscale Coupling of Architecture and Biointerfaces Drives Osteogenesis in 3D-Printed Gyroid Scaffolds

**DOI:** 10.64898/2026.04.16.718992

**Authors:** Pratheesh Kanakarajan Vijaya Kumari, Jasmine Carpenter, Barnett Cleon, Christopher J Panebianco, Joel D Boerckel, Derrick Dean, Vineeth M Vijayan

## Abstract

Engineering functional bone scaffolds can be enhanced by integrating biologically instructive nanoscale surface features (e.g., nanotopography and nanoroughness), micro-scale geometric cues (e.g., curvature and porosity), and macro-scale mechanical properties (e.g., bulk stiffness); however, these length scales are often optimized independently. Here, we present a multiscale design framework combining additive manufacturing of triply periodic minimal surface (TPMS) gyroid scaffolds with plasma-assisted nanoscale surface engineering to regulate osteogenesis. Controlled variation in strut thickness generates distinct architectural regimes with coupled changes in curvature, porosity, and compressive modulus, recapitulating key aspects of trabecular bone mechanics. Micro-computed tomography confirms trabecular bone-like features, while finite element modeling and compression testing reveal that thinner architectures (0.6 mm) exhibit curvature-preserving geometry and distributed stress profiles favorable for cellular interaction. A low-temperature plasma electroless reduction (PER) strategy enables controlled silver nanoparticle deposition, while polydopamine-mediated adhesion ensures uniform and cytocompatible coatings. Notably, PDA-AgNP-functionalized 0.6 mm scaffolds significantly outperform unmodified and AgNP-only groups, exhibiting enhanced cytoskeletal organization, stress fiber formation, matrix mineralization, and osteogenic gene expression. These findings demonstrate that coupling nanoscale biointerface features with micro- and macro-scale architecture produces a synergistic enhancement in osteogenesis, providing a design framework for functional bone scaffolds.

**Table of Content Graphics:** A plasma-enabled strategy integrates 3D-printed scaffold architecture with nanoscale surface engineering to enhance bone formation. By combining tunable structural design with uniform nanoparticle coating, the study shows that optimal biological responses occur only when mechanical and surface cues act together, highlighting a synergistic multiscale approach for designing advanced biomaterials for bone regeneration.

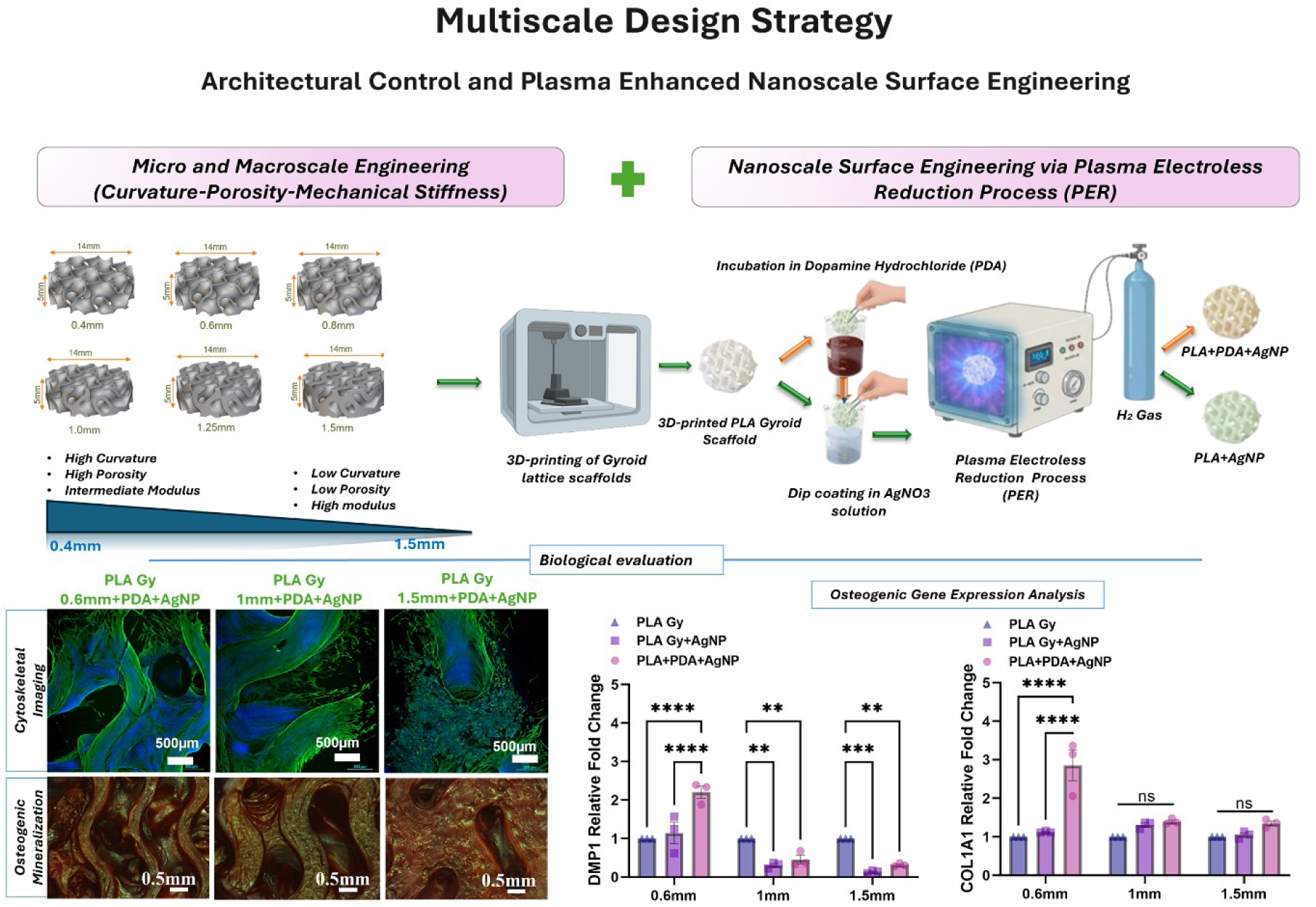

## 1. Introduction

Bone defects arising from trauma, degenerative diseases, or tumor resection represent a significant global burden, with an estimated 2.2 million cases reported annually [1][2]. Although autologous bone grafting remains the clinical gold standard [3], its application is limited by donor site morbidity and restricted availability [4] thereby driving the demand for synthetic bone graft substitutes and engineered scaffolds [5]. Native bone exhibits a hierarchical organization spanning nanoscale collagen mineral composites, microscale trabecular curvature and porosity, and macroscale load-bearing architectures [6] Successful bone regeneration therefore requires coordinated regulation across these structural length scales [7].

Considerable progress in bone tissue engineering has focused on architected scaffolds with controlled geometry, porosity, and mechanical stiffness [8]. Triply periodic minimal surface (TPMS) gyroid structures have emerged as promising trabecular bone analogues due to their interconnected pore networks, high surface-to-volume ratios, and tunable mechanical properties [9][10]. Numerous studies have demonstrated that TPMS-derived architectural mechanics, such as curvature and stiffness, can influence osteogenic differentiation through mechanotransductive pathways [11]. However, most existing approaches primarily optimize bulk architectural parameters while treating surface bioactivity as an independent modification. In reality, these features are not biologically independent, as cells simultaneously sense mechanical cues arising from scaffold architecture and nanoscale features at the cell–material interface. Architectural parameters such as curvature and stiffness regulate cytoskeletal organization and force generation, while nanoscale surface features influence local cell adhesion and contact guidance. Together, these cues converge at the level of cytoskeletal remodeling and downstream osteogenic signaling. As a result, coordinated optimization of both architectural mechanics and nanoscale biointerface properties is required, yet this integrated, multiscale design approach remains insufficiently explored.

To address hierarchical design requirements, several strategies have combined additive manufacturing with electrospinning [12], biomimetic mineral coatings [13], or nanoparticle incorporation to introduce nanoscale features onto 3D-printed scaffolds [14]. While such hybrid systems enhance surface complexity, they often suffer from limited penetration into complex three-dimensional lattices [15], multistep fabrication processes [16], inconsistent interfacial stability, and challenges in achieving uniform nanoscale functionalization across intricate architectures [17]. These limitations underscore the need for scalable and controllable approaches that enable stable nanoscale surface modulation within architecturally complex scaffolds.

Among nanoscale bioactive agents, silver nanoparticles (AgNPs) have attracted considerable interest due to their broad-spectrum antimicrobial properties [18] and reported ability to modulate osteogenic signaling pathways through reactive oxygen species (ROS)-mediated mechanisms and downstream transcriptional activation [19]. At controlled concentrations, AgNPs have been shown to enhance osteogenic differentiation [20]; however, excessive nanoparticle deposition and uncontrolled ion release may induce oxidative stress and cytotoxicity [21]. Consequently, AgNP-functionalized scaffolds have been predominantly developed for antimicrobial protection [14], while their osteogenic potential remains underexplored [22]. Conventional nanoparticle synthesis and anchoring strategies including wet chemical reduction, electrostatic adsorption, and thermal decomposition are typically multistep processes that provide limited control over nanoparticle distribution, stability, and release kinetics, particularly on complex three-dimensional architectures [23]. These challenges necessitate efficient surface engineering strategies capable of stabilizing nanoscale silver biointerfaces while mitigating cytotoxic effects.

Plasma electroless reduction (PER) provides a scalable platform for in situ synthesis and deposition of metallic nanoparticles, such as gold and silver, under low-temperature hydrogen plasma conditions, enabling uniform nucleation without chemical reducing agents [24]. Previous work from our group has demonstrated the ability of PER to immobilize metallic nanoparticles stably on polymeric substrates with controlled surface density and minimal processing complexity [24,25]. While such stable immobilization is essential, effective translation of metallic nanoparticles for tissue engineering applications requires careful modulation of the cell–material biointerface, particularly to regulate nanoparticle stability, protein adsorption, and potential oxidative stress arising from ion release. Polydopamine (PDA), a bioadhesive coating rich in catechol and amine functional groups, enhances nanoparticle anchoring while improving surface hydrophilicity and protein adsorption [26] and has been reported to exhibit antioxidant-like behavior that may buffer reactive oxygen species [27]. The synergistic integration of PDA pre-coating with PER-mediated AgNP deposition therefore presents a rational strategy to stabilize nanoscale silver biointerfaces, mitigate cytotoxicity concerns, and modulate cellular responses within architecturally defined scaffolds.

In this study, a plasma-enabled multiscale design framework is developed that integrates architecturally defined mechanical regimes in TPMS gyroid scaffolds with nanoscale biointerface modulation via PER-deposited AgNPs, with and without PDA pre-coating. Strut thickness is systematically varied to generate distinct micro-scale curvature–porosity landscapes and macro-scale stiffness profiles. The PER process is then employed to achieve uniform nanoparticle deposition across these geometries, and the stabilizing role of PDA is evaluated. Mechanical performance is assessed through finite element analysis and compressive testing, while biological responses are characterized through cytoskeletal organization, mineralization, and osteogenic gene expression. To the best of current knowledge, this represents the first demonstration of plasma-enabled nanoscale biointerface engineering coupled with architecturally defined TPMS mechanical regimes to modulate osteogenesis within a unified and scalable multiscale framework.

## 2. Results and Discussion

### 2.1. Architecturally Defined Mechanical Regimes in TPMS Gyroid Scaffolds

To establish an architecture-driven mechanical baseline prior to nanoscale biointerface modulation, we systematically varied strut thickness to generate distinct mechanical microenvironments within a constant TPMS gyroid topology. Because surface curvature, porosity, and bulk stiffness collectively influence mechanotransduction [28–30], we designed these variations to produce geometry-defined mechanical regimes. These regimes enable differential regulation of cellular responses through architecture-driven mechanical cues.

Given the known sensitivity of osteogenic responses to such mechanically defined microenvironments [28][29], identifying an architecturally optimized condition was therefore essential before introducing nanoscale surface cues. Gyroid triply periodic minimal surface (TPMS) scaffolds with strut thicknesses ranging from 0.4 to 1.5 mm were designed using nTopology and fabricated via extrusion-based 3D printing (Fig. 1A). Printing fidelity was preserved across all geometries, including thin struts (0.4–0.6 mm), with no observable structural defects. The original gyroid topology was maintained across thicknesses, enabling controlled evaluation of geometry-driven mechanical effects.

**Figure 1:**
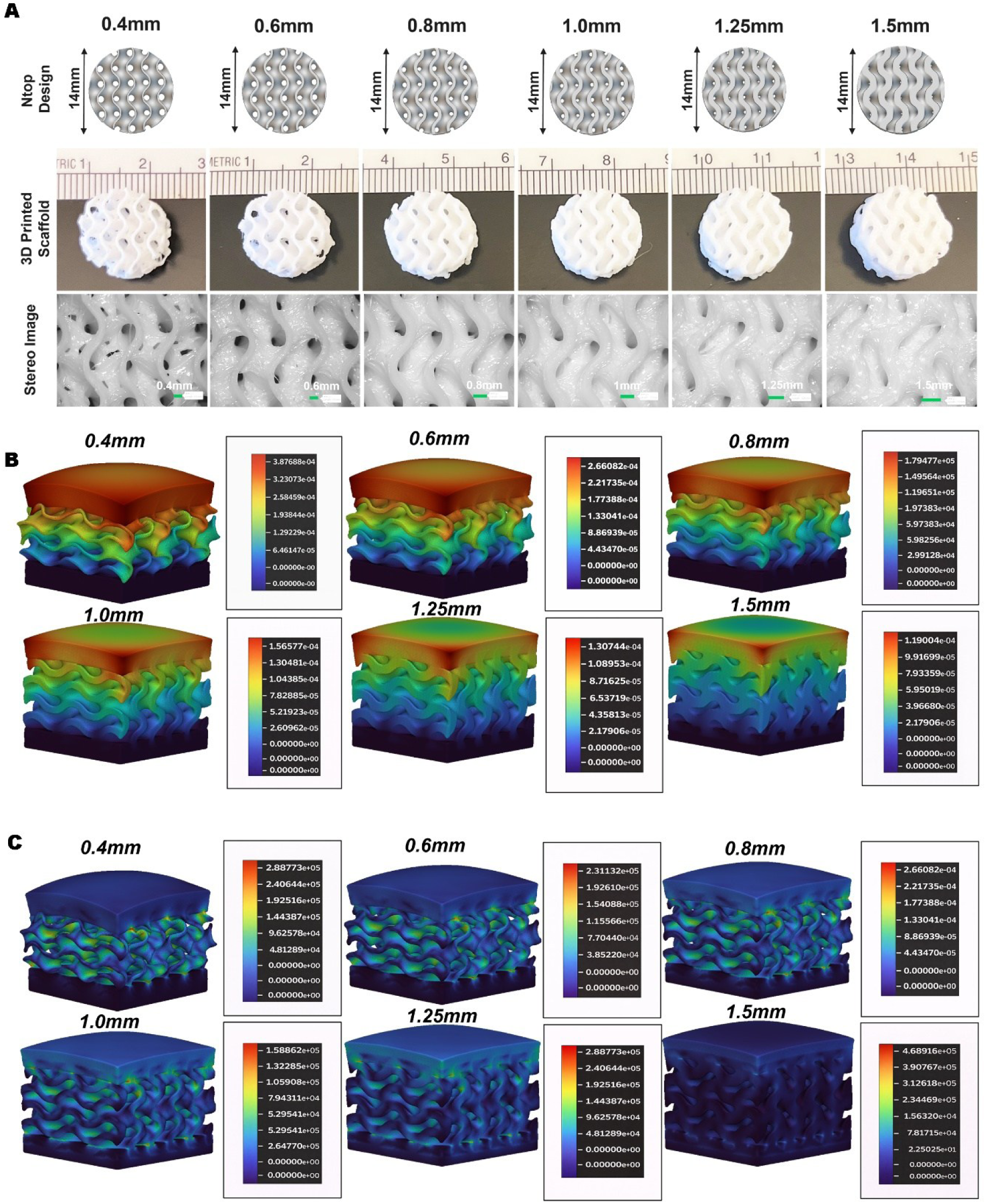
(A) TPMS gyroid scaffold designs with strut thicknesses of 0.4–1.5 mm, corresponding photographs of 3D-printed PLA scaffolds, and stereomicroscopic images showing printed surface morphology. All scaffolds were fabricated with a constant diameter and height of 14 mm. (B) Finite element analysis (FEA)-derived displacement distribution of TPMS gyroid scaffolds with increasing strut thickness under uniaxial compression. (C) Corresponding FEA-derived von Mises stress distribution, illustrating thickness-dependent mechanical response under identical loading conditions.

Systematic variation in strut thickness generated distinct architectural regimes characterized by coupled changes in curvature, porosity, and bulk stiffness. Thinner scaffolds (0.4–0.6 mm) exhibited higher surface curvature and more open pore networks, whereas thicker scaffolds (1.25–1.5 mm) displayed reduced pore openness and increasingly dense architecture. Importantly, the gyroid topology remained intact across all thicknesses, ensuring that mechanical differences arose from controlled geometric modulation rather than topological distortion [29][30].

Finite element analysis (FEA) revealed thickness-dependent deformation behavior under uniaxial compression (Fig. 1B). Thinner scaffolds (0.4–0.6 mm) exhibited greater overall deformation with pronounced displacement gradients across the structure, consistent with more compliant, curvature-preserving architectures typical of trabecular bone [32]. Intermediate thicknesses (0.8–1.0 mm) demonstrated more moderate deformation, suggesting a transition toward more balanced load transfer within the network. In contrast, thicker scaffolds (1.25–1.5 mm) showed reduced displacement, reflecting increased stiffness and a shift toward bulk-dominated mechanical behavior.

Analysis of von Mises stress distributions (Fig. 1C and Table S2) further quantified thickness-dependent load-bearing characteristics. Mean von Mises stress decreased progressively from 9.69 × 10⁴ Pa (0.4 mm) to 1.60 × 10⁴ Pa (1.5 mm), indicating a reduction in overall stress magnitude. Similarly, maximum stress values decreased from 1.49 × 10⁵ Pa to 4.21 × 10⁴ Pa with increasing thickness, suggesting reduced peak stress intensity. However, spatial variability in stress distribution remained substantial across all groups, as reflected by standard deviation (SD) values (5.19 × 10⁴ Pa to 1.45 × 10⁴ Pa) and maximum-to-mean stress ratios (Table S2), indicating heterogeneous load distribution within the architectures. Notably, thinner scaffolds exhibited more localized stress concentrations at nodal junctions and curved strut regions, consistent with junction-dominated deformation governed by architectural curvature, as observed in TPMS-based structures [31][32]. In contrast, thicker scaffolds displayed lower absolute stress magnitudes but retained heterogeneous stress distributions, reflecting a transition toward stiffness-dominated load-bearing behavior. Together, these results indicate that strut thickness modulates both the magnitude and spatial distribution of mechanical cues within the scaffold while preserving the underlying gyroid topology [31][32].

To confirm that systematic modulation of strut thickness generated distinct architectural environments spanning micro-scale geometric cues and macro-scale mechanical behavior, curvature analysis and mechanical validation were performed (Fig. 2A–D). Mean Gaussian curvature remained relatively stable (∼ −1.39 to −1.29 mm⁻²) across scaffolds from 0.4 to 1.0 mm, indicating preservation of the saddle-like negative curvature intrinsic to gyroid minimal surfaces (Fig. 2A) [31]. Beyond 1.0 mm, the curvature progressively deviated, with measurable reduction in surface complexity at 1.25 and 1.5 mm. This indicates that thin and intermediate scaffolds preserve micro-scale geometric heterogeneity, whereas thicker scaffolds progressively lose curvature-driven surface features as strut bulk increases [32].

**Figure 2:**
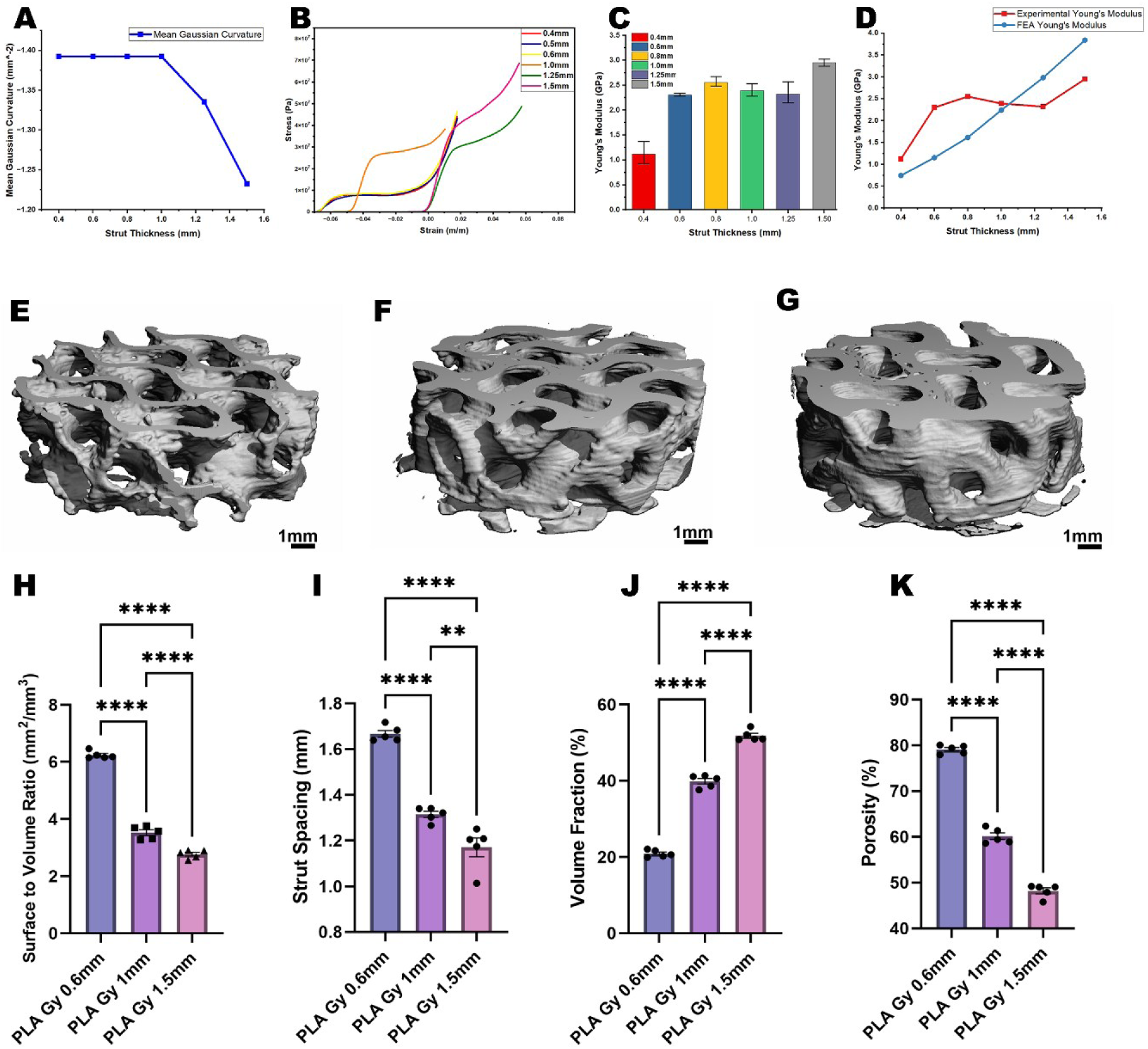
Mechanical characterization on PLA gyroid scaffolds with varying strut thickness. A) Mean Gaussian curvature (mm⁻²) as a function of strut thickness, demonstrating the geometric tunability of the gyroid architecture. B) Representative compressive stress–strain curves for scaffolds across a range of strut thicknesses (0.4–1.6 mm). C) Experimentally determined Young’s modulus (GPa) as a function of strut thickness. D) Comparison of experimental and finite element analysis (FEA)-predicted Young’s modulus values across strut thicknesses, confirming the predictive accuracy of the computational model. Representative micro-computed tomography (micro-CT) 3D reconstructions of PLA gyroid scaffolds with sturt thickeness of E) 0.6 mm, F) 1 mm, and G) 1.5 mm (Scale bars = 1 mm). Quantitative morphological analysis of H) surface-to-volume ratio (mm²/mm³), I) strut spacing (mm), J) volume fraction (%), and K) porosity (%) across the three scaffold designs (PLA Gy 0.6 mm, PLA Gy 1 mm, and PLA Gy 1.5 mm). Data are presented as mean ± SEM (*n* = 5). Statistical significance was determined by one-way ANOVA with Tukey’s post hoc test (** *p* < 0.01, **** *p* < 0.0001).

Mechanical testing further revealed thickness-dependent macro-scale load-bearing behavior. Representative stress–strain curves (Fig. 2B) demonstrated that thin scaffolds (0.6 mm) exhibited lower initial stiffness, earlier yielding, and extended plateau regions characteristics of bending-dominated, trabecular-like mechanics [33]. Intermediate scaffolds (0.8–1.0 mm) displayed a three-phase response with balanced elastic deformation and energy dissipation. In contrast, thicker scaffolds (1.25–1.5 mm) showed higher peak stresses and reduced plateau regions, consistent with increased stiffness and a transition toward bulk-dominated compressive behaviour.

Experimental Young’s modulus values (Fig. 2C) increased from 0.4 mm to 0.8 mm, plateaued between 0.8–1.25 mm, and rose again at 1.5 mm. This nonlinear trend suggests a shift from geometry-dominated mechanics in thin scaffolds to mixed geometry–material behavior in intermediate scaffolds, and ultimately toward stiffness-dominated bulk mechanics at 1.5 mm [30]. FEA predictions showed a more linear increase with thickness (Fig. 2D), while experimental values exhibited deviations likely attributable to printing-induced surface irregularities and microstructural imperfections. Importantly, the 0.6 mm scaffolds exhibited modulus values within the reported range of native trabecular bone, reinforcing their biomechanical relevance [33].

Together, curvature preservation, stress distribution patterns, and compressive modulus measurements define three mechanically distinct architectural regimes: curvature-preserving junction-dominated (0.6 mm), transitional balanced (1.0 mm), and stiffness-driven bulk-dominated (1.5 mm). These three representative conditions span a broad architectural spectrum and were therefore selected to evaluate architecture-dependent biological responses.

### 2.2. Micro- and Macro Scale Architectural Validation of TPMS Gyroid Scaffolds

Based on the mechanically distinct regimes identified from finite element analysis (FEM) of the six initial scaffold designs, three representative strut thicknesses 0.6 mm, 1.0 mm and 1.5 mm were selected for micro-computed tomography (micro-CT) analysis. These thicknesses span the range from curvature-preserving to stiffness-dominated architectures, enabling evaluation across the full spectrum of architecture-driven mechanical behaviour while minimizing redundancy in downstream characterization.

Micro-CT analysis was used to quantitatively capture key microscale architectural features of TPMS gyroid scaffolds, including surface-to-volume ratio, Strut spacing, volume fraction and porosity which are comparable with the features of native trabecular bone as BS/BV, Tb.Sp, BV/TV, 1-BV/TV respectively [34]. These parameters are widely used to describe trabecular bone microarchitecture and provide insight into the local surface availability, pore interconnectivity, and solid-void balance that govern cell-material interactions.

surface-to-volume ratio decreased significantly with increasing strut thickness (Fig.2D) (*p* < 0.0001), indicating reduced surface area available for cell attachment and biological interaction on thicker scaffolds [33][34]. Since osteogenic processes are strongly surface-mediated, higher surface-to-volume ratio in thinner scaffolds suggests a more favorable cell-interactive microenvironment, like highly active trabecular bone surfaces. Sturt spacing showed a reduction from 0.6 mm to 1.5 mm scaffolds (Fig.2E) (*p* < 0.01), reflecting decreased interpore spacing. While lower Sturt spacing indicates a denser structure, larger spacing in thinner scaffolds is more consistent with interconnected porous networks in trabecular bone, which support cell infiltration and nutrient transport.

The volume fraction increased substantially with strut thickness (p < 0.0001), while porosity decreased proportionally (Fig.2G) (p < 0.0001) [36]. These parameters define the solid-void distribution within the scaffold, where the high porosity (∼79% in 0.6 mm scaffolds) falls within the reported range of trabecular bone porosity (40-95%) [33]. whereas thicker scaffolds approach lower porosity regimes associated with denser bone structures. This comparison indicates that thinner scaffolds more closely replicate the microscale porous architecture of trabecular bone, which is known to support enhanced permeability, cell infiltration, and osteogenic potential. These structural parameters confirm that thinner strut scaffolds (0.6 mm) provide higher surface area and porosity conducive to cellular infiltration and nutrient transport [37], whereas thicker struts progressively reduce these biologically favorable characteristics despite maintaining structural integrity [38][39]. Importantly, when considered collectively, micro-CT parameters define a microscale architectural signature that can be used to predict scaffold performance. The combination of high BS/BV, high porosity, and larger interpore spacing in 0.6 mm scaffolds suggests a microenvironment that is more representative of trabecular bone and therefore more conducive to osteogenic differentiation.

Thus, micro-CT analysis not only validates scaffold architecture but also serves as a predictive tool to assess the suitability of engineered scaffolds for bone regeneration prior to biological evaluation. Furthermore, these findings highlight that strut thickness serves as a key design parameter to systematically tune microscale features such as porosity, surface area, and pore spacing, enabling controlled modulation of scaffold microenvironments relevant to bone tissue engineering.

#### 2.2.1. Cytoskeletal Organization and Quantitative Analysis Across TPMS Architectures

To quantitatively assess the influence of scaffold architecture on cellular organization, fluorescence microscopy combined with image analysis was performed on TPMS gyroid scaffolds with varying strut thicknesses [35][36].

Fluorescence imaging revealed distinct differences in actin organization (Fig. 3A), with 0.6 mm scaffolds exhibiting well-aligned, densely organized actin stress fibers, while 1.0 mm and 1.5 mm scaffolds displayed progressively reduced alignment and diffuse organization (Fig. 3B)[37][38]. Quantitative analysis demonstrated that actin intensity decreased with increasing strut thickness, with 0.6 mm scaffolds showing substantially higher values compared to 1.5 mm scaffolds (**p < 0.01) (Fig. 3C) [39][40]. Interestingly, stress fiber density and orientation coherency remained statistically unchanged across all architectures (Fig. 3D and E) [41], indicating that architectural effects on cytoskeletal organization manifest primarily through alterations in overall actin expression and spatial arrangement rather than fiber quantity or directional coherency [42]. These findings establish that TPMS geometric parameters modulate cellular mechanotransduction capacity through architecture-dependent regulation of actin intensity [43][44], with geometrically optimized 0.6 mm scaffolds promoting the most robust cytoskeletal organization [45]

**Figure 3:**
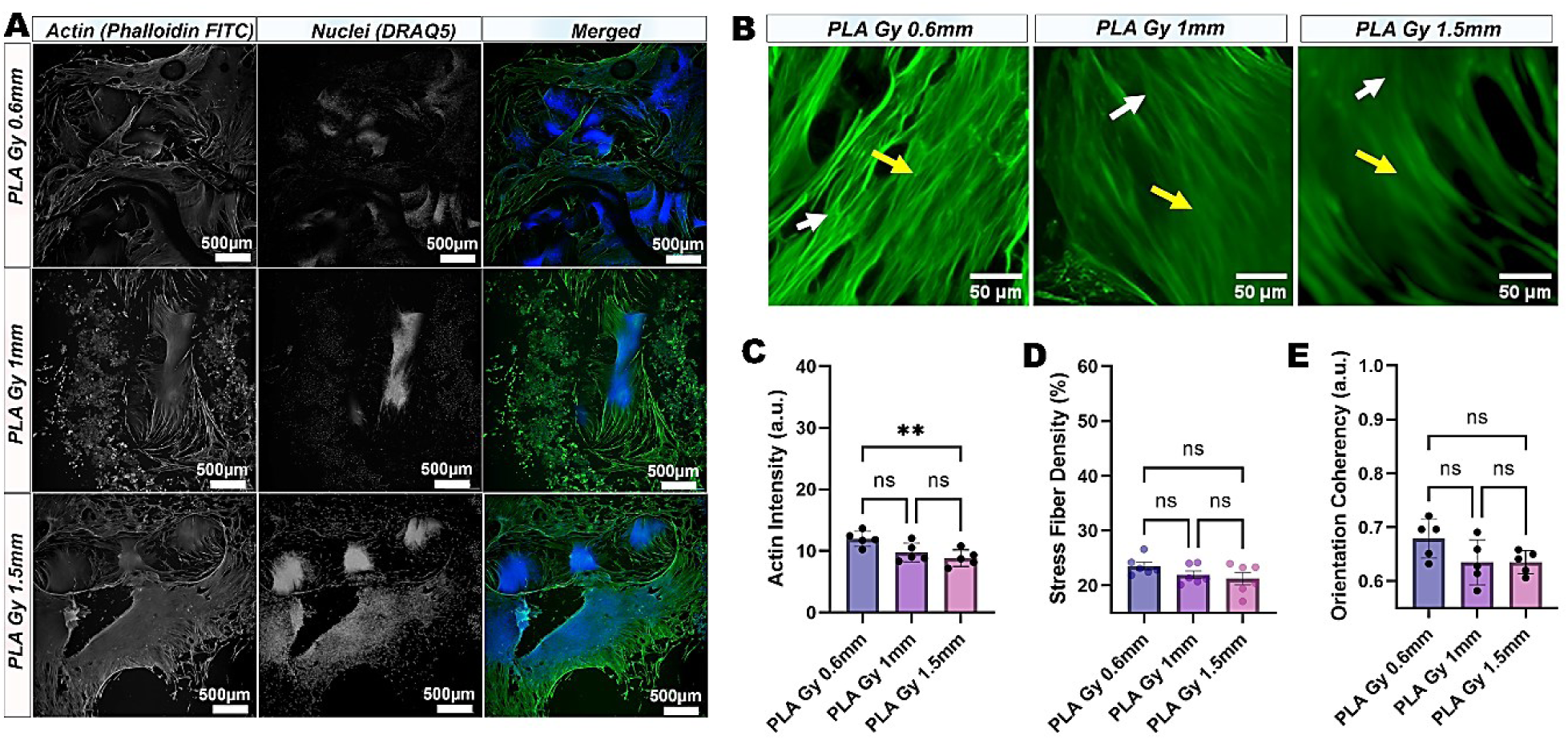
Cytoskeletal response of cells cultured on PLA gyroid scaffolds with varying strut thickness. A) Representative fluorescence micrographs of cells seeded on PLA Gy 0.6 mm, PLA Gy 1 mm, and PLA Gy 1.5 mm scaffolds, stained for filamentous actin (F-actin (Phalloidin-FITC, green), nuclei (DRAQ5, far-red pseudocolored to blue) and their merged overlay. Scale bars = 500 µm. B) High-magnification fluorescence images of actin cytoskeleton organization on each scaffold group (Yellow and white arrows indicate aligned stress fibers and actin bundles respectively. Scale bars = 50-80 µm). Quantitative analysis of C) actin fluorescence intensity (a.u.), D) stress fiber density (%), and E) orientation coherency (a.u.) across scaffold groups. Data are presented as mean ± SEM (*n* = 5). Statistical comparisons were performed using one-way ANOVA with Tukey’s post hoc test (** *p* < 0.01; ns, not significant).

#### 2.2.2. Architecture-Driven Osteogenic Mineralization

The influence of architectural mechanics on osteogenic differentiation was evaluated through Alizarin Red staining [46] at 7, 14, and 21 days (Fig. 4). At the early time point (7 days), minimal mineral deposition was observed across all strut thicknesses, with no significant differences between groups, consistent with the early proliferative phase preceding matrix maturation [47]. By day 14, mineralization increased in all scaffolds; however, the 0.6 mm group exhibited significantly higher calcium deposition compared to the 1.5 mm scaffolds (Fig. 4B). The 1.0 mm scaffolds demonstrated intermediate mineralization levels, suggesting that transitional architectural regimes partially support osteogenic progression but do not maximize mineral deposition [48]. At 21 days, architectural differences became more pronounced. The 0.6 mm scaffolds displayed the highest mineral accumulation, followed by the 1.0 mm group, while 1.5 mm scaffolds exhibited substantially lower mineral deposition (Fig. 4B). These findings indicate that curvature-preserving, junction-dominated mechanical environments enhance late-stage matrix mineralization [49], whereas bulk-dominated stiffness attenuates osteogenic maturation [50].

**Figure 4:**
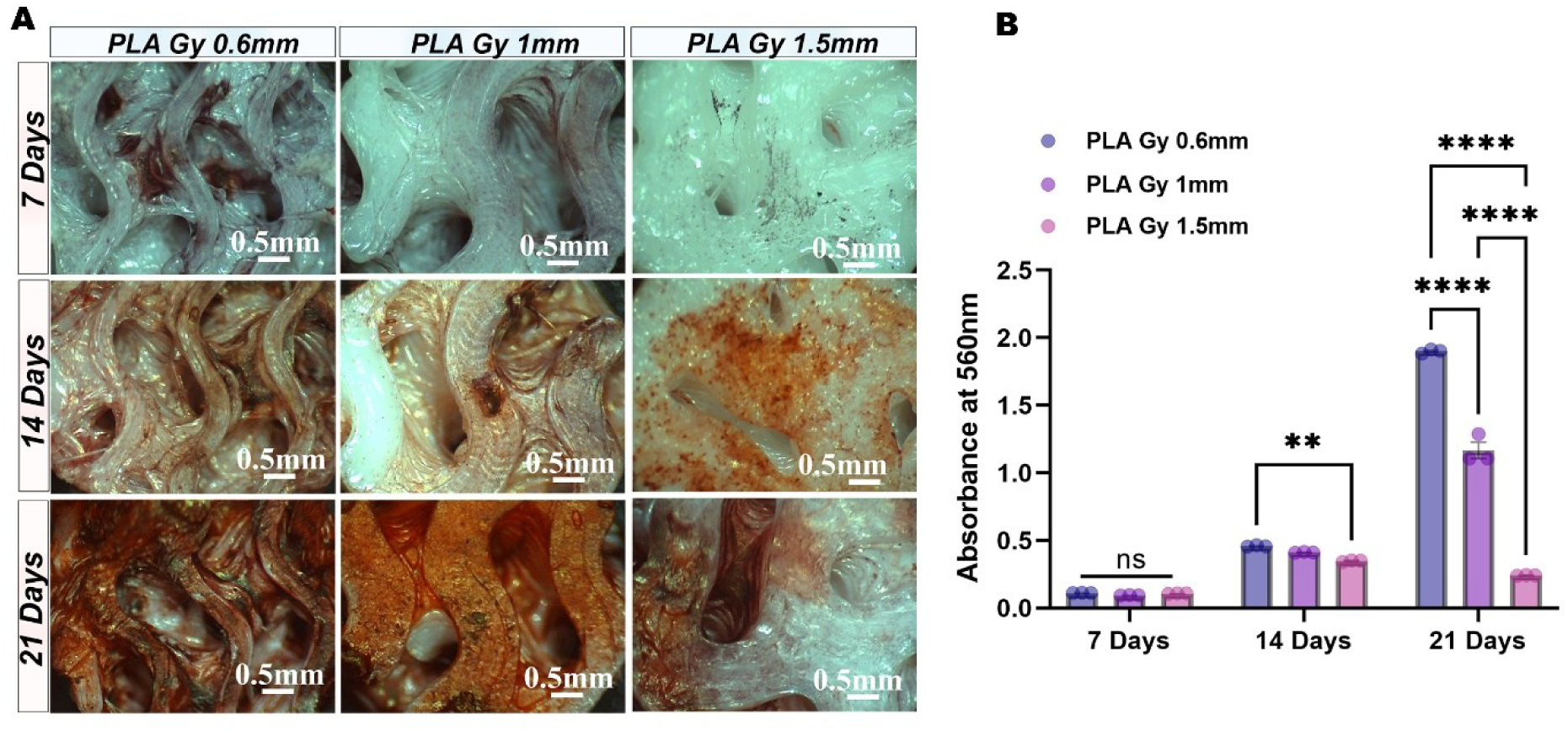
Osteogenic mineralization of cells cultured on PLA gyroid scaffolds over 21 days. A) Representative stereomicroscopy images of PLA Gy 0.6 mm, PLA Gy 1 mm, and PLA Gy 1.5 mm scaffolds at 7, 14, and 21 days under osteogenic culture conditions, stained with Alizarin Red S to visualize calcium deposition. Progressive reddish-orange staining over time indicates increasing extracellular matrix mineralization across all scaffold groups. Scale bars = 0.5 mm. B) Quantitative spectrophotometric analysis of Alizarin Red S absorbance at 560 nm across scaffold groups at 7, 14, and 21 days, reflecting the degree of mineral deposition over the culture period. Data are presented as mean ± SEM (n = 3). Statistical comparisons were performed using two-way ANOVA with Tukey’s post hoc test (** p < 0.01, **** p < 0.0001; ns, not significant).

Together, these results demonstrate that architectural mechanics regulate not only the magnitude of mineral deposition but also the temporal stability of osteogenic progression. The 0.6 mm scaffold provides the most favorable balance of curvature complexity and trabecular-like mechanical compliance [51], while 1.0 mm scaffolds represent an intermediate regime and 1.5 mm scaffolds shift toward stiffness-dominated behavior that dampens osteogenic efficiency [51][52]. These findings justify the selection of 0.6, 1.0, and 1.5 mm scaffolds as representative architectural regimes spanning micro-scale curvature preservation to macro-scale stiffness dominance for subsequent nanoscale surface integration studies.

### 2.3. Plasma-Enabled Nanoscale Biointerface Engineering and Concentration Optimization

#### 2.3.1. Plasma Electroless Reduction Strategy for In Situ AgNP Immobilization

Following identification of three representative architectural regimes (0.6, 1.0, and 1.5 mm), nanoscale surface modification was introduced using plasma electroless reduction (PER). This approach enables in situ reduction of Ag⁺ ions and simultaneous immobilization of silver nanoparticles (AgNPs) directly onto the PLA surface (Fig. 4A), resulting in stable nanoparticle anchoring without the need for external chemical reducing agents or post-deposition stabilization steps [24]. A mild post-treatment sonication step ensures removal of loosely bound species, retaining only firmly immobilized nanoparticles.

In contrast to conventional wet-chemical methods, where nanoparticle synthesis and deposition are typically separate processes, PER integrates these steps into a single plasma-driven process. This in situ reduction mechanism facilitates conformal nanoparticle deposition across complex three-dimensional scaffold geometries while maintaining process simplicity and environmental compatibility [24]. These characteristics make PER particularly well-suited for introducing nanoscale biointerfaces onto architected polymeric scaffolds.

#### 2.3.2. Concentration-Dependent Morphology and Cytocompatibility Screening

To decouple nanoparticle optimization from architectural variables, initial concentration screening was conducted on PLA scaffolds fabricated with a simple honeycomb lattice infill (45% density) (Fig. 5A). This reproducible geometry enabled systematic evaluation of nanoparticle nucleation behavior, surface distribution, and cytocompatibility prior to application onto TPMS gyroid scaffolds. SEM imaging directly on PLA substrates revealed clear concentration-dependent nanoparticle morphology (Fig. 5B). At higher precursor concentrations (25 and 12.5 mM), PLA + AgNP surfaces exhibited clustered and partially agglomerated nanoparticle domains, indicative of dense and less regulated nucleation [53]. As the precursor concentration decreased, nanoparticles became progressively more discrete and spatially separated. At 0.7 mM concentration of AgNO_3_ the nanoparticles appeared uniformly distributed and stably anchored with minimal aggregation. To further enhance nanoparticle immobilization, PLA surfaces were pre-coated with polydopamine (PDA) prior to PER treatment. Even at higher precursor concentrations such as 25 mM, PDA-precoated surfaces exhibited more individually resolved and less clustered nanoparticles compared to non-PDA coated surfaces. This observation suggests that catechol-mediated chelation of Ag⁺ [54] ions improve nucleation control and interfacial stabilization during plasma reduction [55]. Figure S3 shows an increase in the emission intensity consistent with the increase in the concentration of silver nanoparticles for both PLA + AgNP and PLA + PDA + AgNP scaffolds. The PLA + PDA + AgNP exhibited a slightly higher intensity at 15 mM and 25 mM as compared to the PLA + AgNP. While further studies need to be performed, reports have from others have indicated the laser induced breakdown spectroscopy (LIBS) can be enhanced by the presence of nanoparticles. This elevated uniformity in the signal intensity is likely due to the catechol and AgNP nucleation chemistry [56]. The enhancement is dependent on the interparticle separation as well [57].

**Figure 5:**
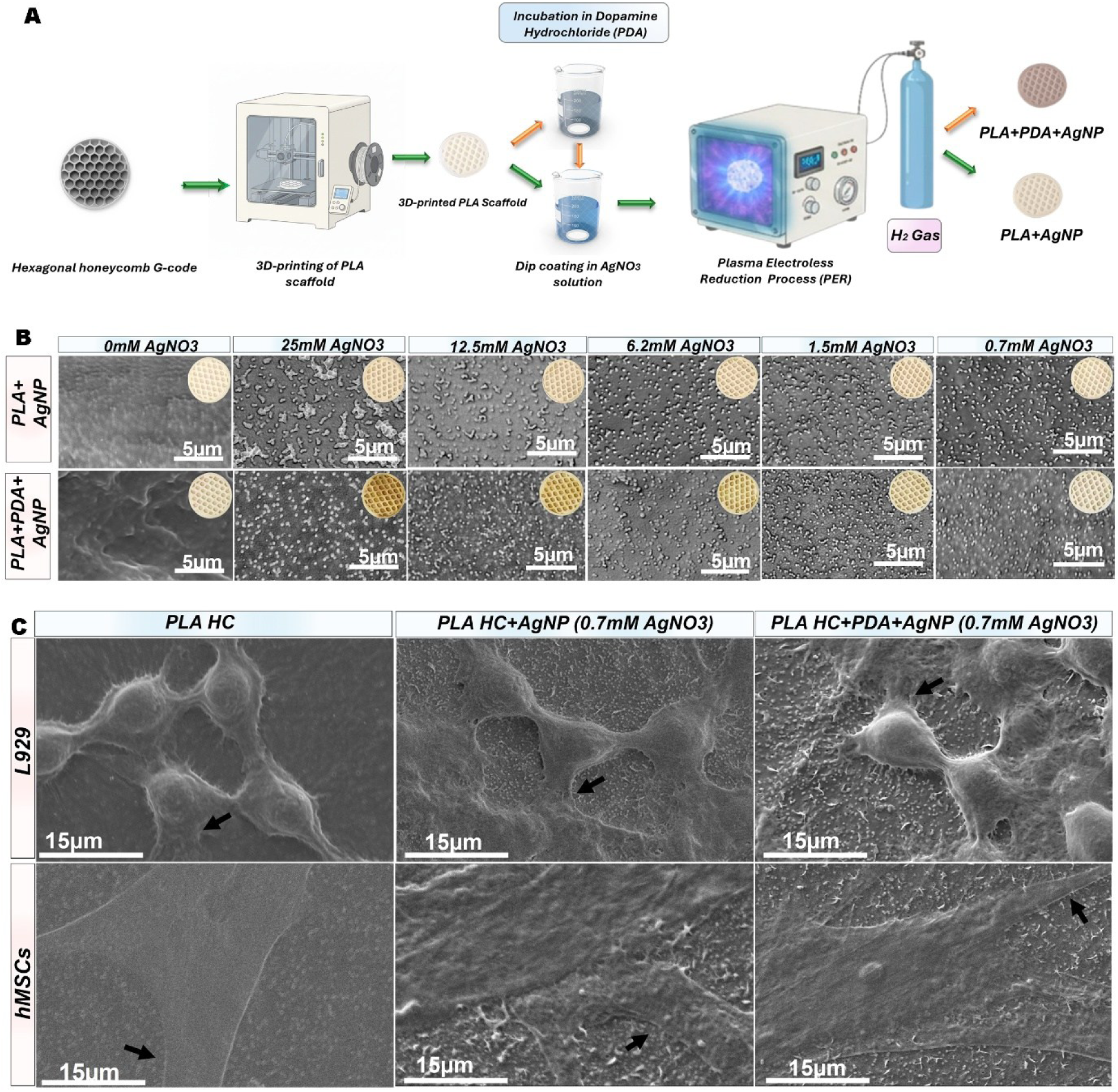
Fabrication and surface characterization of silver nanoparticle (AgNP)-functionalized PLA scaffolds. A) Schematic illustration of the scaffold fabrication and surface modification workflow: hexagonal honeycomb G-code was used to 3D-print PLA scaffolds, which were subsequently dip-coated in AgNO₃ solution, with or without prior incubation in polydopamine hydrochloride (PDA), followed by Plasma Electroless Reduction (PER) under H₂ gas to yield PLA+AgNP and PLA+PDA+AgNP constructs, respectively. B) Representative scanning electron microscopy (SEM) images of PLA+AgNP (top row) and PLA+PDA+AgNP (bottom row) scaffolds fabricated across a range of AgNO₃ concentrations (0–25 mM), with corresponding optical images of the scaffold surface shown as insets. Scale bars = 5 µm. C) SEM micrographs of L929 fibroblasts (top row) and human mesenchymal stem cells (hMSCs, bottom row) adhered to unmodified PLA HC, PLA HC+AgNP (0.7 mM AgNO₃), and PLA HC+PDA+AgNP (0.7 mM AgNO₃) scaffolds. Black arrows indicate representative cell–scaffold interactions. Scale bars = 15 µm.

Collectively, these findings highlight that nanoparticle formation via PER is governed by a balance between precursor concentration–driven nucleation and interfacial stabilization. While higher concentrations promote increased nanoparticle density, they also introduce aggregation and heterogeneity that may compromise surface uniformity. In contrast, intermediate concentrations, particularly around 0.7 mM, provide an optimal regime for achieving uniformly distributed and stably anchored nanoparticles. The incorporation of PDA further refines this process by enhancing nucleation control and surface adhesion, resulting in a more consistent nanoscale interface. This controlled nanoparticle architecture establishes a well-defined and reproducible biointerface, which is critical for isolating the role of nanoscale cues in subsequent cell–material interaction studies on TPMS gyroid scaffolds.

TEM imaging (Fig. S4) provided an approximate estimation of nanoparticle size and morphology. Because nanoparticles were deposited onto conductive copper grids rather than PLA substrates for TEM analysis, these images do not directly represent anchoring behavior on polymer surfaces. Instead, TEM confirms general particle distribution trends and average size of 110 ± 20 nm on uncoated and 135 ± 30 nm on PDA coated (Fig S4). Consistent with SEM observations, nanoparticles formed without PDA exhibited broader size heterogeneity and aggregation, whereas PDA-associated nanoparticles appeared more discrete and relatively uniform. At 24 hours, the Alamar Blue assay demonstrated consistently high metabolic activity across all scaffold groups (Fig. S5)., indicating good cytocompatibility of PLA, PDA coated, and AgNP modified scaffolds. No statistically significant differences were observed among the groups, suggesting the AgNP concentrations have not induced acute cytotoxic effects within the initial culture period. Biological SEM imaging further confirmed stable cell attachment and distinct differences in cell morphology across on 0.7 mM-modified surfaces. The upper panel (L929 fibroblasts) and lower panel (hMSCs) (Fig. 5C) demonstrate that both cell types exhibited limited spreading with relatively smooth contours on the pristine PLA scaffolds. In contrast, AgNP-modified surfaces promoted increased surface interaction, while PDA-assisted AgNP coatings further enhanced cell anchorage and spreading. Notably, cells on PLA HC+PDA+AgNP displayed extensive cytoplasmic extensions and flattened morphology (arrows), forming intimate contact with the underlying substrate (Fig 5C). These broad protrusive structures are characteristic of mesenchymal stem cell spreading and suggest enhanced adhesion and mechanosensitive interactions with the nano-engineered surface, supporting a surface-driven mechanotransductive response [58]. PDA-modified groups exhibited broader spreading and more continuous surface interaction relative to AgNP-only surfaces. No morphological evidence of acute cytotoxicity was observed at the optimized concentration. Collectively, concentration-dependent morphology and cytocompatibility screening identified 0.7 mM AgNO₃ as the optimal precursor concentration. This condition was therefore selected for application onto the architecturally defined TPMS gyroid scaffolds to investigate cross-scale coupling.

### 2.4. Cross-Scale Coupling of Architectural Mechanics and Nanoscale Bioactivity

#### 2.4.1. Architecture Surface Interplay Governs Cytoskeletal Organization

Following independent optimization of architectural mechanics (0.6, 1.0, and 1.5 mm) and nanoparticle concentration (0.7 mM AgNO₃), the three mechanically distinct TPMS gyroid regimes were functionalized with AgNP via PER, with and without PDA pre-coating (Fig. 6A). This sequential strategy ensured that nanoscale biointerface engineering was introduced within well-defined curvature-dependent mechanical microenvironments. To quantitatively assess cytoskeletal responses to scaffold architecture and surface modifications, fluorescence microscopy combined with image analysis was performed across all conditions (Fig. 6 and 7). We hypothesized that nanoscale biointerface engineering would not independently dictate cellular behaviour but rather amplify osteogenic mechanotransduction preferentially within curvature-preserving architectural regimes [59][60].

**Figure 6:**
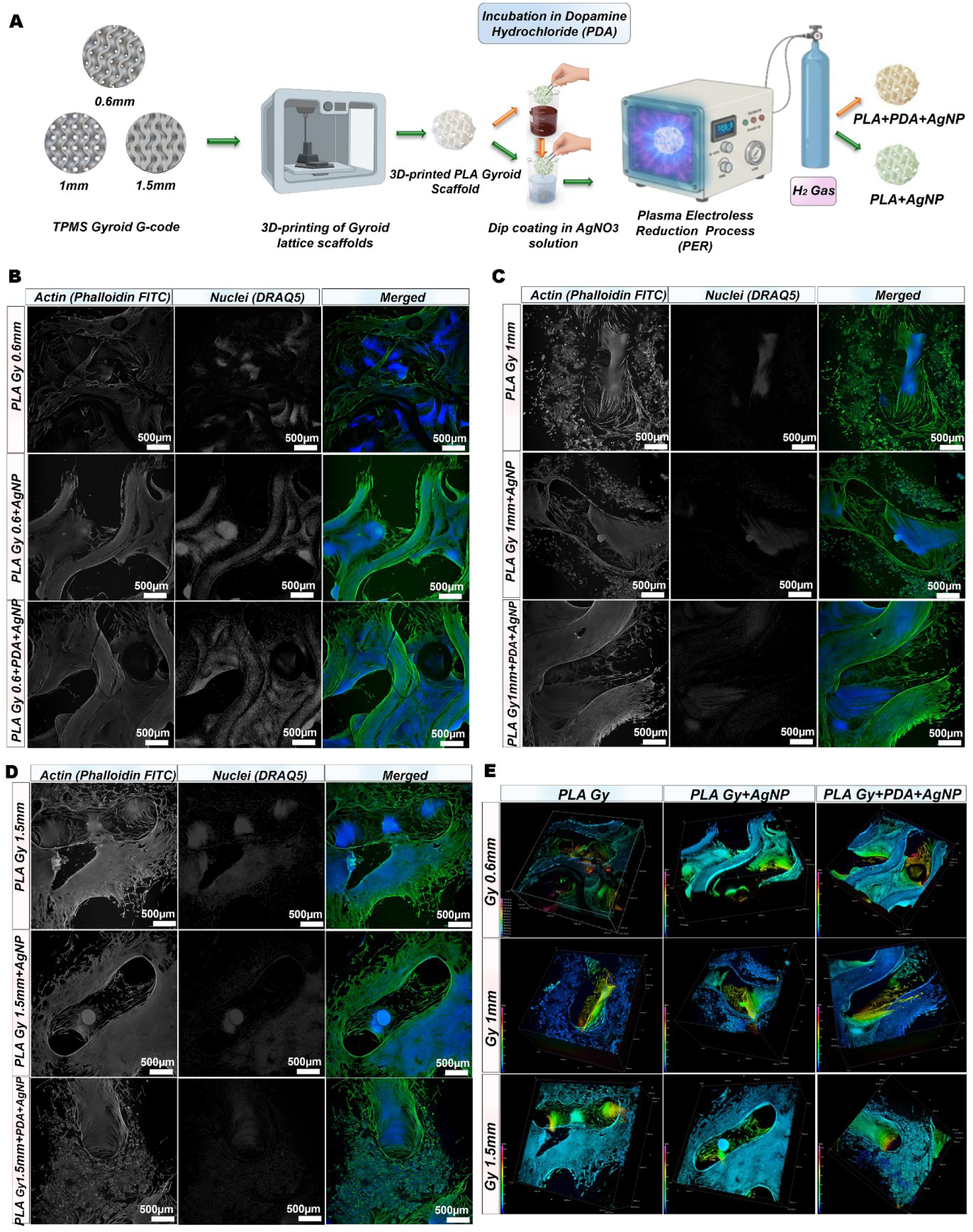
Cytoskeletal organization of cells cultured on AgNP-functionalized PLA gyroid scaffolds with varying unit cell sizes. A) Schematic illustration of the gyroid scaffold fabrication and surface modification workflow: TPMS gyroid G-codes at unit cell sizes of 0.6 mm, 1 mm, and 1.5 mm were used to 3D-print PLA gyroid scaffolds, which were subsequently dip- coated in AgNO₃ solution with or without prior polydopamine (PDA) incubation, followed by Plasma Electroless Reduction (PER) under H₂ gas to produce PLA+AgNP and PLA+PDA+AgNP constructs. Representative fluorescence micrographs of cells seeded on unmodified and AgNP-functionalized scaffolds at unit cell sizes of B) 0.6 mm, C) 1 mm, and D) 1.5 mm, stained for filamentous actin (F-actin (Phalloidin-FITC, green), nuclei (DRAQ5, far-red pseudocolored to blue), and their merged overlay. For each unit cell size, images are shown for PLA Gy (unmodified), PLA Gy+AgNP, and PLA Gy+PDA+AgNP groups. Scale bars = 500 µm. E) High magnification confocal fluorescence images of the merged actin and nuclear channels, highlighting differences in cytoskeletal morphology and spatial organization across scaffold surface treatments (PLA Gy, PLA Gy+AgNP, and PLA Gy+PDA+AgNP) for each unit cell size (0.6 mm, 1 mm, and 1.5 mm).

**Figure 7:**
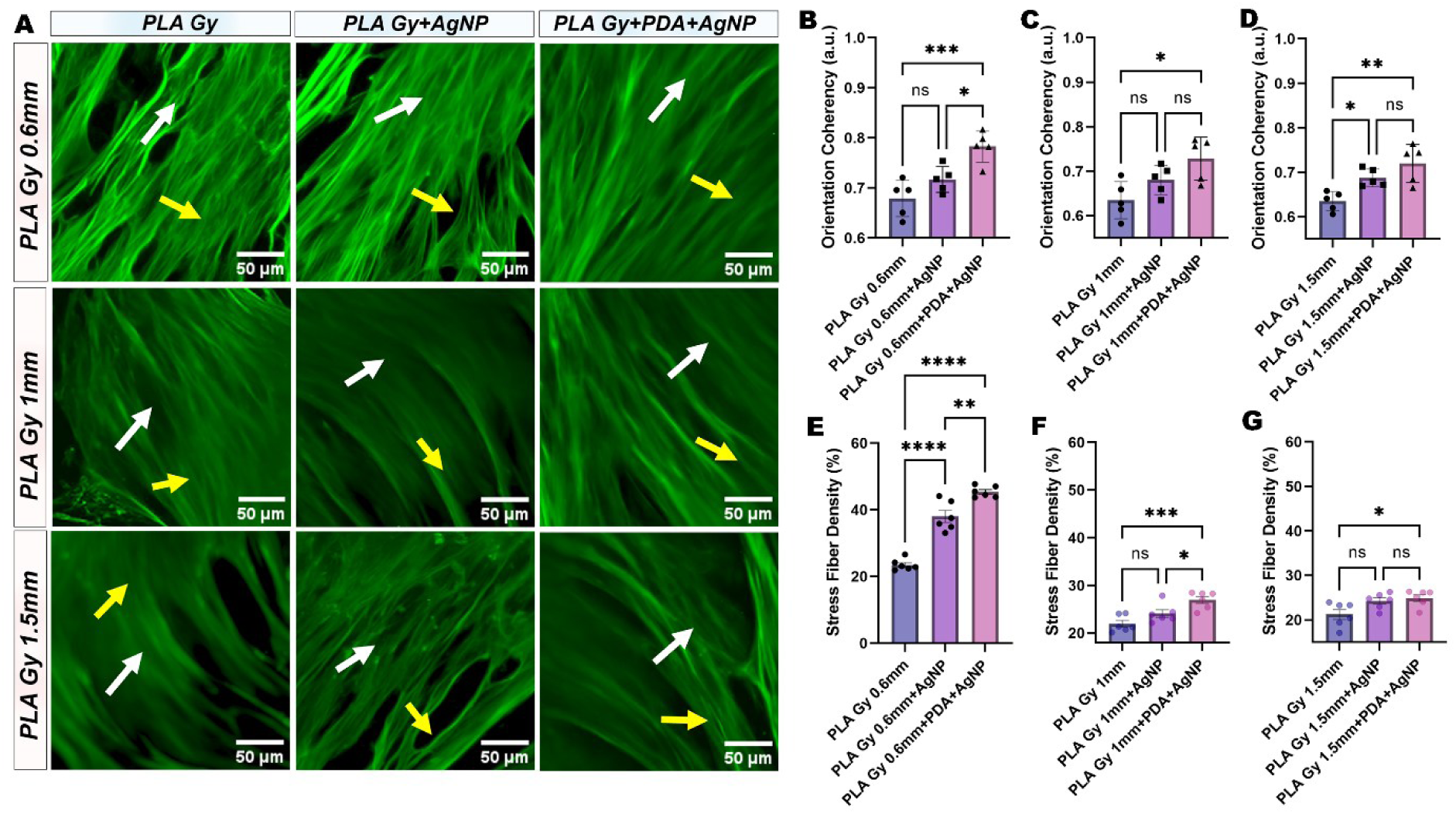
Actin cytoskeleton organization of cells cultured on PLA scaffolds with and without PDA and silver nanoparticle surface modifications. A) Representative fluorescence microscopy images of phalloidin-stained actin filaments in cells seeded on PLA Gyroid scaffolds with 0.6 mm, 1 mm, and 1.5 mm, either unmodified (PLA Gy), functionalized with silver nanoparticles (PLA Gy+AgNP) and coated with polydopamine prior to silver nanoparticle deposition (PLA Gy+PDA+AgNP). Scale bars = 50 µm. B–D) Quantification of actin intensity (a.u.) for strut periodicities of 0.6 mm, 1 mm, and 1.5 mm, respectively. E–G) Stress fiber density (white arrows) across the three different t scaffold strut thicknesses. H–J) Orientation coherency (yellow arrows.) as a measure of actin alignment directionality for on 0.6 mm, 1 mm, and 1.5 mm, respectively. Data are presented as mean ± SEM (n = 5). Statistical significance was performed by one-way ANOVA with Tukey’s post hoc test: *p < 0.05, **p < 0.01, ***p < 0.001, ****p < 0.0001; ns = not significant.

Specifically, we postulated that stabilized nanoscale adhesion cues would reinforce cytoskeletal organization most effectively in mechanically permissive microenvironments, while stiffness-dominated architectures would limit such amplification [60]. Three-dimensional depth reconstruction (Fig. 6E) revealed maximal cellular infiltration within the curvature network in 0.6 mm + PDA + AgNP scaffolds, whereas thicker scaffolds exhibited greater surface localization. These findings indicate that optimal cytoskeletal integration requires synergy between accessible curvature geometry and stabilized nanoscale adhesion chemistry.

Surface modification altered cytoskeletal organization in an architecture-dependent manner (Fig. 6B-E and Fig.7). Actin intensity analysis demonstrated robust surface modification effects across all strut thicknesses, with PDA+AgNP scaffolds exhibiting significantly higher values compared to AgNP-only and pristine PLA (****p < 0.0001) (Fig. S6). Notably, 0.6 mm scaffolds showed the highest absolute actin intensity Fig 7B) (30 a.u.), progressively declining to 15-20 a.u. on 1.5 mm scaffolds, indicating strong architecture dependence. Stress fiber density followed similar trends, with significant differences on 0.6 mm scaffolds (****p < 0.0001) that diminished on thicker architectures, becoming predominantly non-significant on 1.5 mm scaffolds (Fig. 7E-G) [39]. Interestingly, orientation coherency remained unchanged across conditions (Fig. 7H-J), suggesting that surface modifications primarily enhance actin expression and polymerization rather than inducing fiber reorganization [61] [43]. Together, these results demonstrate that PDA-assisted AgNP functionalization enhances cytoskeletal organization, with effects most pronounced in curvature-preserving architectures were surface chemistry and geometry act synergistically.

#### 2.4.2. Geometry-Dependent Amplification of Osteogenesis

Mineralization analysis demonstrated that nanoscale functionalization enhances osteogenic differentiation in a geometry-dependent manner (Fig. 8A–F). At 7 days, mineral deposition was minimal across all groups. By 14 days, 0.6 mm scaffolds functionalized with PDA+AgNP exhibited significantly greater calcium deposition than PLA and AgNP-only controls, indicating that nanoscale bioactivity most effectively accelerates osteogenesis within curvature-preserving mechanical environments [62][63]. At 21 days, a hierarchical pattern emerged. In 0.6 mm scaffolds, PDA+AgNP yielded the highest mineralization, followed by AgNP alone and PLA controls. In 1.0 mm scaffolds, surface mineralization was relative to PLA, yet absolute levels remained below the optimized 0.6 mm condition. In 1.5 mm scaffolds, AgNP and PDA+AgNP coatings partially enhanced osteogenesis compared to unmodified PLA. However, mineral deposition remained inferior to curvature-preserving architectures [62].

**Figure 8:**
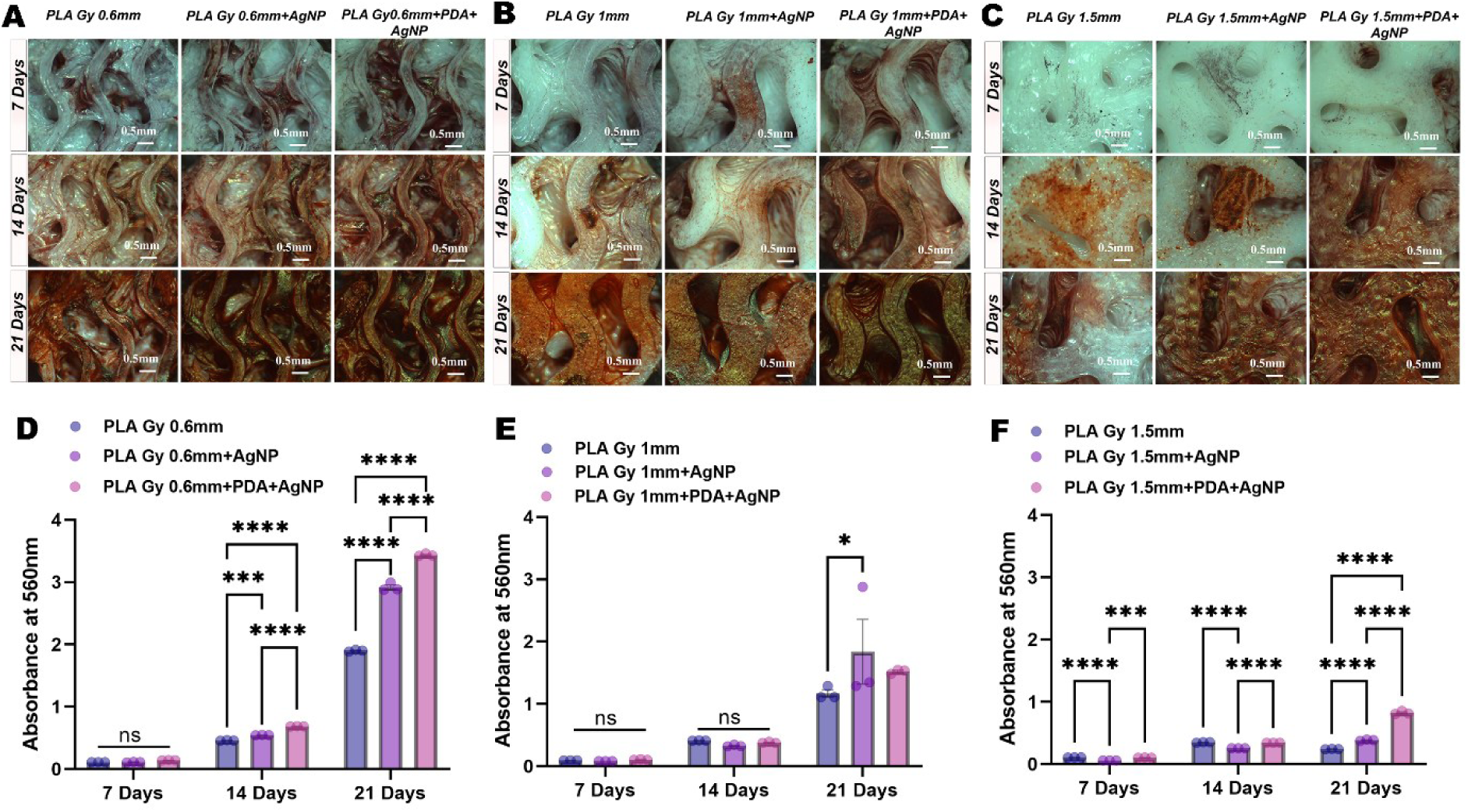
Osteogenic mineralization of cells cultured on AgNP-functionalized PLA gyroid scaffolds over 21 days. Representative stereomicroscopy images of Alizarin Red S-stained scaffolds at 7, 14, and 21 days for A) PLA Gy 0.6 mm, B) PLA Gy 1 mm, and C) PLA Gy 1.5 mm groups, comparing unmodified (PLA Gy), silver nanoparticle-functionalized (PLA Gy+AgNP), and polydopamine-mediated silver nanoparticle-functionalized (PLA Gy+PDA+AgNP) scaffolds. Progressive reddish-orange staining over time reflects increasing extracellular matrix mineralization across all groups. Scale bars = 0.5 mm. Quantitative spectrophotometric analysis of Alizarin Red S absorbance at 560 nm across time points (7, 14, and 21 days) for D) PLA Gy 0.6 mm, E) PLA Gy 1 mm, and F) PLA Gy 1.5 mm scaffold groups. Data are presented as mean ± SEM (*n* = 3). Statistical comparisons were performed using two-way ANOVA with Tukey’s post hoc test (* *p* < 0.05, *** *p* < 0.001, **** *p* < 0.0001; ns, not significant).

Further the gene expression profiling at 21 days confirmed coordinated architectural surface synergy (Fig. 9A–F). RUNX2 expression was maximal in 0.6 mm PDA+AgNP scaffolds with progressive attenuation at 1.0 mm and 1.5 mm (Fig 9A). Matrix maturation markers DMP1 (Fig 9B), IBSP (Fig 9C), SPP1 (Fig 9D) and late differentiation markers OCN (Fig 9E), COL1A1(Fig 9F) [62][64] supported the similar geometry-dependent trends. Although nanoscale functionalization enhanced expression relative to unmodified PLA within each thickness group, the magnitude of upregulation remained architecture-limited [62].

**Figure 9:**
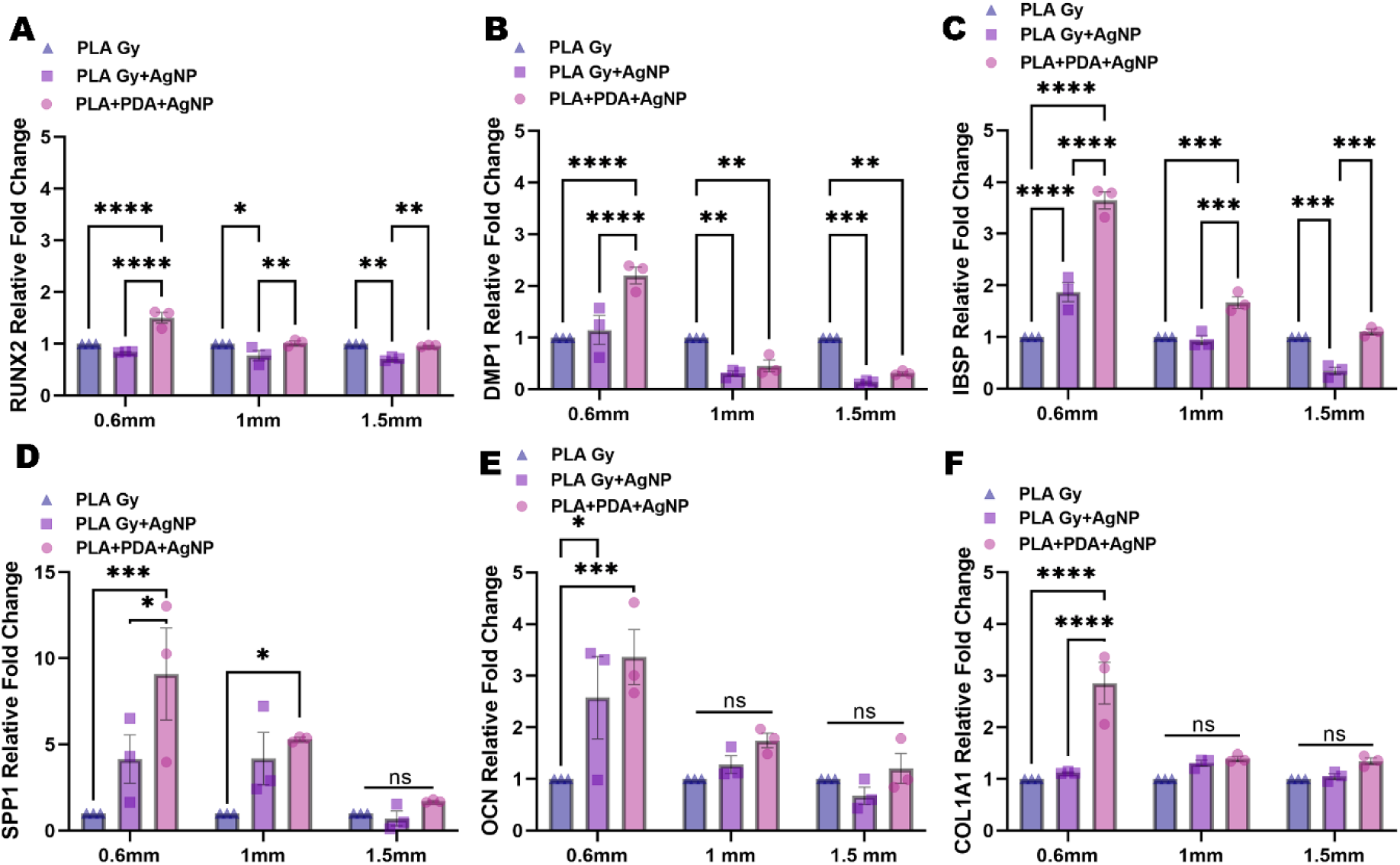
Osteogenic gene expression in cells cultured on AgNP-functionalized PLA gyroid scaffolds. Relative fold change in mRNA expression of osteogenic markers A) RUNX2, B) DMP1, C) IBSP, D) SPP1, E) OCN, and F) COL1A1, quantified by RT-qPCR in cells cultured on PLA Gy, PLA Gy+AgNP, and PLA Gy+PDA+AgNP scaffolds across unit cell sizes of 0.6 mm, 1 mm, and 1.5 mm. Gene expression is normalized to the housekeeping gene and presented as mean ± SEM (*n* = 3). Statistical comparisons were performed using two-way ANOVA with Tukey’s post hoc test (* *p* < 0.05, ** *p* < 0.01, *** *p* < 0.001, **** *p* < 0.0001; ns, not significant).

Collectively, these results support a hierarchical cross-scale coupling model in which architectural mechanics establish the primary mechanobiological framework [65], while nanoscale biointerface engineering tunes signal transmission efficiency [64]. The curvature-preserved 0.6 mm scaffolds, enhanced cytoskeletal alignment likely facilitates effective mechanotransductive signalling, enabling PDA-stabilized AgNP interfaces to amplify osteogenic transcriptional programs [64]. In contrast, stiffness-dominated 1.5 mm scaffolds limit osteogenic amplification despite nanoscale modification. The concordance between cytoskeletal organization and RUNX2 upregulation is consistent with tension-dependent pathways such as RhoA/ROCK and YAP/TAZ signaling [66], although these were not directly interrogated.

This multiscale coupling concept is summarized schematically in Fig. 10, which illustrates the progressive enhancement of osteogenic mineralization from stiffness-dominated to curvature-preserving architectures, and its further amplification through nanoscale PDA-AgNP surface functionalization.

**Figure 10:**
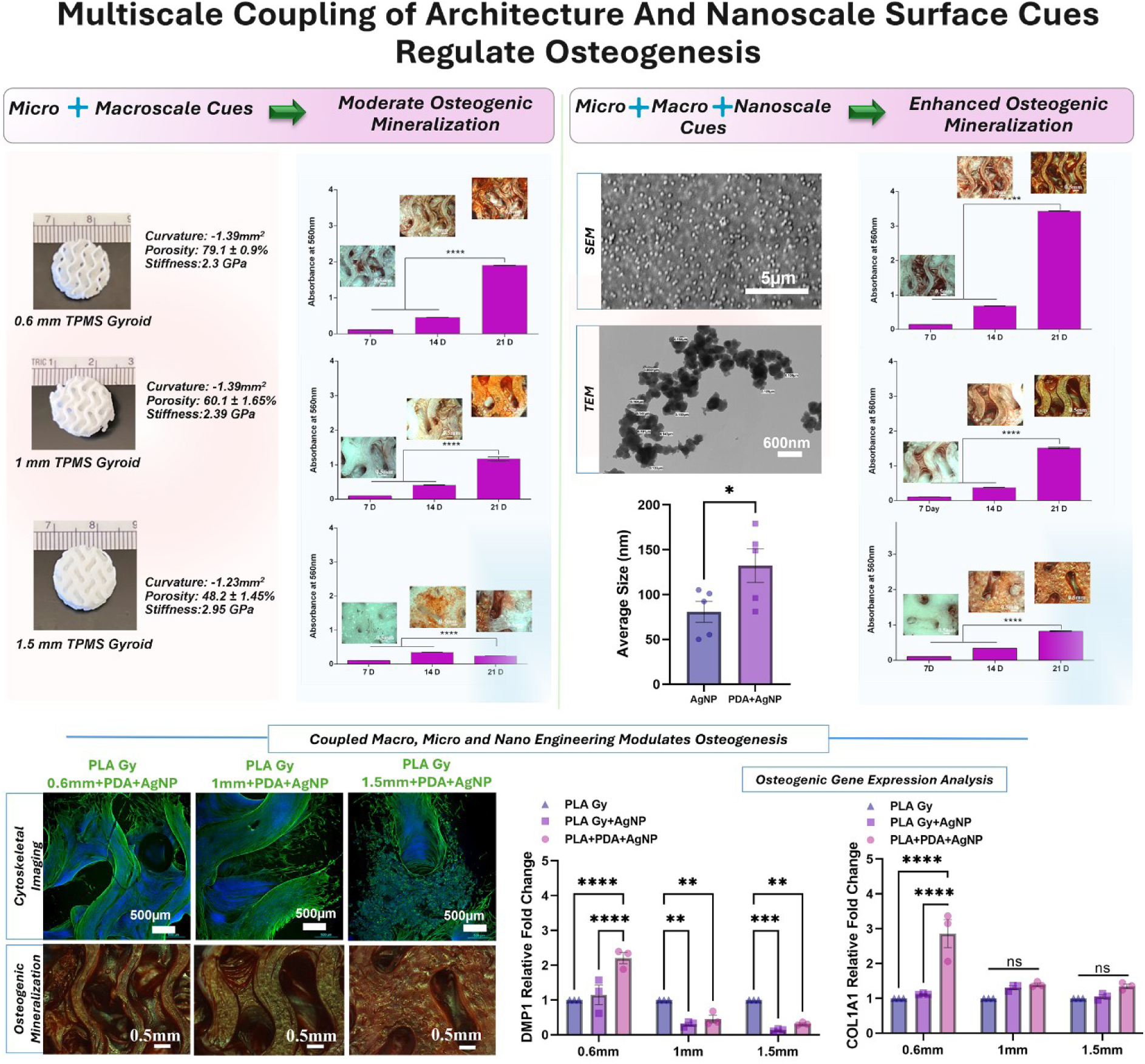
Schematic overview of multiscale coupling between architectural and nanoscale surface cues in regulating osteogenesis. TPMS gyroid scaffolds with varying strut thickness establish distinct mechanical microenvironments, yielding moderate osteogenic responses when micro- and macroscale cues act independently. Incorporation of plasma-enabled PDA– AgNP biointerfaces enhances mineralization across all conditions; however, maximal osteogenesis is achieved in curvature-preserving scaffolds (0.6 mm), where nanoscale cues synergistically amplify architecture-driven cellular responses. These results establish a key design principle: coordinated coupling of micro-, macro-, and nanoscale cues is required to achieve optimal osteogenic performance.

## 3. Conclusions

This study establishes a hierarchical cross-scale design framework in which architectural mechanics define the primary mechanobiological environment and nanoscale surface engineering tunes signal amplification rather than acting independently. Through systematic modulation of TPMS gyroid strut thickness, we identified three mechanically distinct regimes spanning curvature-preserving, transitional, and stiffness-dominated architectures. Preservation of negative Gaussian curvature and trabecular-like mechanical compliance in 0.6 mm scaffolds proved critical for maintaining elevated actin intensity, coordinated stress fiber distribution, and high orientation coherence, thereby driving sustained cytoskeletal organization and osteogenic differentiation. Independent optimization of plasma electroless reduction (PER) enabled green, in situ deposition of silver nanoparticles directly onto polymer scaffolds, with 0.7 mM AgNO₃ identified as the optimal concentration based on uniform surface distribution and cytocompatibility. Polydopamine (PDA) pre-coating enhanced nanoparticle immobilization, reduced agglomeration, and stabilized the biointerface, thereby improving cell–material interactions without compromising viability. When integrated, architectural and nanoscale cues acted synergistically. PDA-assisted AgNP functionalization amplified osteogenesis most effectively within curvature-preserving 0.6 mm scaffolds, producing the highest mineral deposition and upregulation of RUNX2, DMP1, IBSP, SPP1, OCN, and COL1A1. In contrast, stiffness-dominated 1.5 mm scaffolds exhibited attenuated cytoskeletal integration and reduced osteogenic responsiveness, even after nanoscale modification, demonstrating that surface bioactivity cannot fully override unfavourable macro-scale mechanics. Collectively, these findings demonstrate that computationally defined geometric mechanics can be directly translated into functional biological outcomes when coupled with optimized nanoscale bio interfaces. Rather than treating architecture and surface chemistry as independent variables, this work provides evidence for hierarchical mechanical biochemical coupling as a guiding principle for rational scaffold design. Future investigations should evaluate this cross-scale framework under dynamic loading and in preclinical models to assess translational relevance in load bearing and patient-specific bone regeneration strategies.

## 4. Experimental Section

### 4.1. Fabrication of TPMS Gyroid Lattice Scaffolds via Fused Deposition Modeling 3D Printing

Triply periodic minimal surface (TPMS) gyroid lattice scaffolds were designed using nTopology software (nTopology Inc., USA). Cylindrical scaffold geometries with dimensions of 14 mm in diameter and 5 mm in height were used as the base structure. A gyroid infill pattern with a unit cell size of 8 mm and a tolerance of 0.3 mm was applied to generate the lattice architecture. Strut thickness was systematically varied from 0.4 to 1.5 mm to produce distinct architectural regimes with controlled curvature, porosity, and mechanical properties, while maintaining a constant underlying gyroid topology across all designs.

The finalized designs were exported as STL files and processed using FlashPrint 5 software for slicing. Prior to printing, slicing parameters such as layer height and infill resolution were adjusted to ensure accurate reproduction of the designed lattice features. Scaffolds were fabricated using a fused deposition modeling (FDM) 3D printer (Creator Max, FlashForge, Zhejiang, China) with polylactic acid (PLA) filament (1.75 mm diameter). Printing was performed at a nozzle temperature of 200 °C and a build plate temperature of 50 °C, with a print speed of 30 mm/s and a travel speed of 60 mm/s. These parameters were selected to ensure consistent printing fidelity, dimensional accuracy, and structural integrity across all scaffold designs.

### 4.2 Plasma Electroless Reduction (PER) for Silver Nanoparticle Surface Functionalization of Gyroid Scaffolds

Silver nanoparticle (AgNP) deposition was achieved using a plasma electroless reduction (PER) process adapted from our previously reported protocol [24], with modifications to enable integration with polydopamine (PDA)-mediated surface stabilization. Prior to nanoparticle deposition, 3D-printed PLA gyroid scaffolds were subjected to air plasma activation (13.56 MHz radiofrequency, 45 W, 20 sccm air flow) for 10 min to enhance surface hydrophilicity and promote metal ion adsorption. The activated scaffolds were then immersed in aqueous silver nitrate (AgNO₃) solutions of varying concentrations (0–25 mM) to facilitate adsorption of Ag⁺ ions onto the scaffold surface.

Following ion adsorption, the scaffolds were transferred to a plasma chamber (Harrick Plasma, PDC-001-HP) and exposed to hydrogen plasma (13.56 MHz RF, 45 W, 40 sccm H₂ flow) for 1 min, consistent with the PER framework described in [24]. This plasma treatment induced in situ reduction of surface-bound silver ions, resulting in the formation of uniformly distributed AgNPs across the complex 3D scaffold architecture. The AgNO₃ concentration was systematically varied to control nanoparticle density while minimizing aggregation and preserving scaffold integrity.

To remove loosely bound nanoparticles and ensure coating stability, AgNP-functionalized scaffolds were rinsed with Milli-Q water and subjected to bath sonication (Branson 2510, Danbury, CT, USA) for 5 min, followed by washing in phosphate-buffered saline (PBS, pH 7.4). This post-treatment step ensured retention of stably adhered nanoparticles suitable for subsequent biological evaluation.

For enhanced nanoparticle adhesion and cytocompatibility, a subset of scaffolds was pre-coated with polydopamine (PDA), representing a key modification to the previously reported PER process [24]. Briefly, plasma-activated scaffolds were incubated in a 2% dopamine hydrochloride solution (Thermo Fisher Scientific, Waltham, MA, USA) at 37 °C for 2.5 h, allowing oxidative polymerization to form a conformal PDA layer. AgNP deposition via the PER process was then performed on PDA-coated scaffolds under identical plasma conditions. The PDA layer facilitated uniform nanoparticle anchoring and improved coating stability without altering the underlying scaffold architecture.

To systematically evaluate the effect of nanoparticle density and surface stabilization, two parallel surface modification strategies were implemented across a range of AgNO₃ concentrations. In the first approach, plasma-activated scaffolds were directly immersed in AgNO₃ solutions followed by PER processing to generate AgNP-functionalized surfaces (PLA+AgNP). In the second approach, scaffolds were first pre-coated with PDA and subsequently subjected to the same AgNO₃ immersion and PER process (PLA+PDA+AgNP). This design enabled direct comparison of nanoparticle formation and distribution with and without PDA-mediated adhesion under identical plasma conditions, allowing identification of optimal nanoparticle concentration and coating stability. Based on these comparisons, an intermediate AgNO₃ concentration (0.7 mM) was selected for subsequent biological studies due to its uniform nanoparticle distribution and cytocompatibility.

### 4.3. Finite element analysis (FEA)

Finite element analysis (FEA) was performed using nTopology to evaluate the mechanical behavior of the gyroid scaffolds under compressive loading. The analysis was conducted on gyroid structures with varying strut thicknesses of 0.4mm, 0.6mm, 0.8mm, 1mm, 1.25mm, and 1.5mm. The scaffolds were modeled as isotropic linear elastic materials with material properties corresponding to PLA, including Young’s modulus of 3.5 GPa and Poisson’s ratio of 0.36. A static compression analysis was performed with displacement restraint and force as boundary conditions. A compressive force vector of (0-5000) N was applied to simulate physiological loading conditions. Finite element mesh was generated with appropriate mesh density, and the entity of nodes was defined for the analysis. The simulation evaluated key mechanical parameters including stress distribution, strain, and displacement within the scaffold structures. This analysis enabled the comparison of mechanical performance across different strut thicknesses to optimize scaffold design.

### 4.4. Microcomputed Tomography (Micro-CT) of TPMS Gyroid Scaffolds

High resolution three-dimensional microcomputed tomography (Micro CT) images of the TPMS gyroids were captured in the air with the Micro CT 45 system (SCANCO Medical Ag, Brüttisellen, Switzerland) The imaging parameters applied were: isotropic voxel size of 20 μm, integration time of 300 ms, X-ray intensity of 145 μA, and peak tube voltage of 55 kVp. The PLA scaffold was segmented from the surrounding air using a threshold of -94 mgHA/cm3. For noise suppression a three-dimensional gaussian filter of 3.7 with filter support of 4 was used. The volume fraction, strut and pore parameters were quantified using the manufacturer provided software. Parameters are indicated by standard nomenclature for trabecular bone analysis, but applies to scaffold properties (e.g., “bone volume fraction, BV/TV” refers to scaffold volume fraction).

### 4.5. Fluorescence Imaging of Cytoskeletal Organization in hMSCs on Gyroid Scaffolds

Actin staining was performed to visualize the cytoskeletal organization and cell morphology of hMSCs seeded on the scaffolds according to manufacturer’s protocol. Briefly, hMSCs purchased from ATCC (ATCC, USA: Cat no: PCS-500-012) were seeded on gyroid scaffolds with three different strut thicknesses (0.6mm, 1mm, and 1.5mm), as well as on AgNP-coated and PDA+AgNP-coated gyroid scaffolds and cultured for 48 hours in a CO_2_ incubator (Thermo Scientific, Thermo Fisher Scientific, USA: Model: Forma Series). The scaffolds were then washed gently with 1× Phosphate-Buffered Saline (PBS) (Gibco, Thermo Fisher Scientific, USA: Cat no: 14190-144) in a shaker for three times (3 minutes each) to remove the culture media and fixed with 4% paraformaldehyde (Thermo Fisher Scientific, USA: Cat no: J19943-K2) (PFA) for 45 minutes at room temperature. Following fixation, the scaffolds were washed three times with 1x PBS and permeabilized with 0.1% Triton X-100 (Fisher Scientific, USA: Cat no: BP151-500) in PBS for 15 minutes at room temperature. The permeabilized scaffolds were then incubated with Alexa Fluor 488 Phalloidin in 1% Bovine serum albumin (Thermo Scientific, Thermo Fisher Scientific, USA: Cat no: 37520) (BSA) diluted in PBS at a ratio of 1:400 for 1.5 hours at room temperature to stain the F-actin filaments. Subsequently, nuclear staining was performed using DRAQ5 (Invitrogen, Thermo Fisher Scientific, USA: Cat no: 65088092) at a dilution of 1:2000 (10μM in PBS) for 15 minutes in the dark. The scaffolds were washed thoroughly with 1xPBS to remove unbound dye and imaged using a Nikon confocal laser scanning microscope (Nikon Eclipse Ti2). Images were acquired at 4× and 10× magnifications to visualize cell spreading, morphology, and cytoskeletal organization on the different scaffold surfaces. Z-stack confocal imaging was performed to assess cellular infiltration and distribution within the scaffold architecture and curvature.

### 4.6. Quantitative Analysis of Actin Cytoskeletal Organization and Alignment

Actin cytoskeletal organization was assessed by fluorescence imaging of F-actin stained with phalloidin and imaged using confocal microscopy (Nikon Eclipse Ti2) under identical acquisition settings across all samples. Image analysis was performed using Fiji (ImageJ). Raw images were converted to 8-bit grayscale and fixed-size regions of interest (ROIs; 200x200 pixels per side) were applied in representative cell-covered regions, excluding pores, edges, and acellular areas. Actin fluorescence intensity was quantified as the mean gray value following background subtraction. Stress fiber density was determined by thresholding the actin channel using a consistent algorithm across all samples and calculating the area fraction of filamentous structures within each ROI. Cytoskeletal alignment was evaluated using the OrientationJ plugin based on structure tensor analysis, and orientation coherency values ranging from 0 (random organization) to 1 (perfect alignment) were extracted. Data are presented as mean ± standard error mean (SEM).

### 4.7. Alizarin Red Staining and Quantification of Osteogenic Mineralization

Alizarin red staining was performed on the hard scaffolds on 7, 14 and 21 days to evaluate the mineralization potential. Briefly, Bone Marrow derived Mesenchymal Stem Cells (ATCC) were seeded on the scaffolds at a seeding density of 0.7x105 cells per scaffold. The cells were incubated in mesenchymal stem cell complete media (Basal media: ATCC, USA: Cat no: PCS-500-030 and growth kit: ATCC, USA: Cat no: PCS-500-041) for five days. Subsequently the scaffolds were washed gently with 1xPBS to remove any stem cell media and supplemented with complete osteogenic differentiation media (StemPro™ Osteogenesis Differentiation Kit: Gibco, Thermo Fisher Scientific, USA: Cat no: A1007201). The osteogenic media change was performed every three days. At 7, 14 and 21 day timepoint the scaffolds were gently washed with 1x PBS for three times to remove any residual medium. The scaffolds were then fixed for 45 minutes with 4% PFA at room temperature . The scaffolds were then washed thoroughly with 1x PBS and incubated with 2% alizarin red s solution (MilliporeSigma, USA: Cat no: TMS008C) for 45 minutes with gentle shaking at room temperature. The scaffolds were then washed carefully with PBS to remove the unbound dye. Subsequently the scaffolds were dried overnight at room temperature. Stereo microscopic images of the scaffolds at 5x magnification were taken using Fisher Scientific stereomicroscope equipped with Motic Cam 2.0. The osteogenic mineralization was then quantified using cetylpyridinium chloride (CPC). Briefly, 10% CPC (MP Biomedicals, USA: Cat no: MP219017780) was prepared in 10mM sodium phosphate buffer (pH:7) and filtered using a 0.2µm syringe filter to get the clear solution. The scaffolds were then incubated with 10% CPC for 15 minutes with shaking. Then 50μl elute was transferred to a 96 well plate and optical density was read at 560nm in BioTek Cytation 3 (BioTek, USA) imaging reader.

### 4.8. Scanning electron microscopy (SEM) of the Gyroid Scaffolds

The surface architecture of the scaffolds was examined using a scanning electron microscope (Phenom-XL). The scaffolds were sputter coated with 10nm gold. The scaffolds were then imaged using the backscattered electron detection method. Images were acquired at 30,000x magnification to visualize the surface topography, polydopamine coating and silver nanoparticle distribution.

### 4.9. Biological SEM Analysis of Cell Morphology and Adhesion

Cell morphology and adhesion of L929 cells and human mesenchymal stem cells (hMSCs) seeded on PLA HC, PLA HC + 0.7mM AgNP, and PLA HC + PDA + 0.7mM AgNP honeycomb lattice scaffolds for 48hours were examined using a scanning electron microscope (Phenom-XL). Following culture, scaffolds were gently washed with 1× PBS to remove non-adherent cells and fixed with 2.5% glutaraldehyde in PBS for 2 hours. After rinsing 2-3 times with PBS, samples underwent sequential ethanol dehydration (30%, 50%, 60%, 70%, 80%, 90%, 95%, and 100% twice). The dehydration step was performed for 10 minutes each, followed by overnight immersion in 100% ethanol and air drying. Scaffolds were sputter coated with 5nm gold prior to imaging. Images were captured at 10000x and 20000x magnification.

## 5. Laser-Induced Breakdown Spectroscopy (LIBS) Analysis

Laser-induced breakdown spectroscopy experiments were conducted using a Q-switched neodymium-yttrium aluminium garnet (Nd: YAG) laser from Quantel (Q-Smart 850), operating at the fundamental wavelength of 1064 nm and a repetition rate of 10 Hz. The pulse length was approximately 6 ns, and the energy was adjusted to 200 mJ per pulse using a company-supplied attenuation module. The laser pulse is focused normal to the sample surface using a planoconvex lens with a focal length of 150 mm. All experiments were conducted in air on the same day. Emission from the laser-produced plasma was relay-imaged onto an optical fiber core using a pair of planoconvex lenses of focal length 75 mm. The optical fiber delivered the plasma emission to an 8-channel Avantes spectrometer with a spectral range that covered 200-930 nm. Each spectrum was collected approximately 1.28 ms after plasma formation with an integration time of 1.10 ms. Silver emission was monitored using the Ag I emission line at 546.5 nm. Line intensities were obtained by fitting the emission profile to a Lorentzian function and extracting the area under the curve after baseline correction.

### 5.1. Transmission Electron Microscopy (TEM) Analysis of Silver Nanoparticles

The morphology, size, and distribution of silver nanoparticles were examined using transmission electron microscopy. Copper grids were used as substrates for nanoparticle deposition following the same protocols used for PLA scaffold coating. Briefly, copper grids underwent plasma activation for 10 minutes, followed by dip-coating in 0.7mM AgNO₃ solution and hydrogen plasma treatment for 1 minute to facilitate the reduction reaction. For PDA-coated samples, plasma-activated copper grids were first incubated in 2% dopamine hydrochloride solution for 2.5 hours at 37°C to form a polydopamine coating, followed by the same silver nanoparticle deposition procedure using 0.7mM AgNO₃ solution and hydrogen plasma reduction. The coated copper grids were air-dried at room temperature prior to imaging. The samples were examined using a Tecnai Spirit T12 transmission electron microscope operated at 20-120 kV. Images were acquired at magnifications of 10,000× and 20,000× to analyze the silver nanoparticle size, morphology, and distribution on both uncoated and PDA-coated substrates.

### 5.2. Alamar blue assay for Cell Metabolic Activity

Cell metabolic activity was studied on honeycomb lattice scaffolds and evaluated using the Alamar blue assay based on manufacturer’s protocol on the scaffold extracts prepared following ISO standards (ISO 10993-12) . L929 fibroblast cells were seeded at a density of 1×10⁵ cells per well and cultured for 24 hours. The culture medium was then replaced with scaffold extracts from PLA honeycomb lattice scaffolds and PDA-coated honeycomb lattice scaffolds treated with varying AgNO₃ concentrations (25mM, 12.5mM, 6.2mM, 3.1mM, 1.5mM, and 0.7mM). After 24 hours of incubation at 37°C in a CO₂ incubator, 10% Alamar blue reagent prepared in complete medium was added to each well and incubated for 2 hours at 37°C in a CO₂ incubator. The fluorescence intensity was measured at excitation and emission wavelengths of 560nm and 590nm, respectively, using a BioTek Cytation 3 imaging reader to assess metabolic activity.

### 5.3. Real-time polymerase chain reaction (RT-qPCR) for Osteogenic Gene Expression

Gene expression analysis was performed to evaluate the osteogenic differentiation of hMSCs cultured on the lattice scaffolds. Briefly, hMSCs were seeded on the scaffolds and supplemented with osteogenic differentiation medium for 21 days, with media changes performed every three days. At the 21-day time point, total RNA was extracted from the cells using the TRizol (Invitrogen, Thermo Fisher Scientific, USA: Cat no: 15-596-026) method according to the manufacturer’s protocol. The RNA concentration and purity were quantified using a NanoDrop spectrophotometer (Thermo Scientific, Thermo Fisher Scientific, USA: Model: NanoDrop One). Subsequently, complementary DNA (cDNA) synthesis was performed using the High-Capacity cDNA Reverse Transcription Kit (Catalog number 4368814) following the manufacturer’s instructions. Real-time PCR was conducted using PowerUp™ SYBR™ Green Master Mix for qPCR (Applied Biosystems, Thermo Fisher Scientific, USA: Cat no: A25918) with custom-designed primers for osteogenic marker genes including RUNX2, DMP1, IBSP, SPP1, OCN, and COL1A1 (Table 1). GAPDH was used as the housekeeping gene for normalization. RT-PCR was performed according to manufacturer’s protocol. The PCR cycling conditions consisted of UDG activation at 50°C for 2 minutes, polymerase activation at 95°C for 2 minutes, followed by 40 cycles of denaturation at 95°C for 15 seconds and annealing/extension at 60°C for 1 minute. The reactions were performed using a Bio-Rad real-time PCR system (Bio-Rad Laboratories, USA: Model: CFX Opus 96). Gene expression levels were analyzed using the ΔΔCt method with relative quantification against uncoated bare PLA scaffolds of corresponding strut thicknesses (0.6mm, 1mm, and 1.5mm) as controls.

**Table 1:**
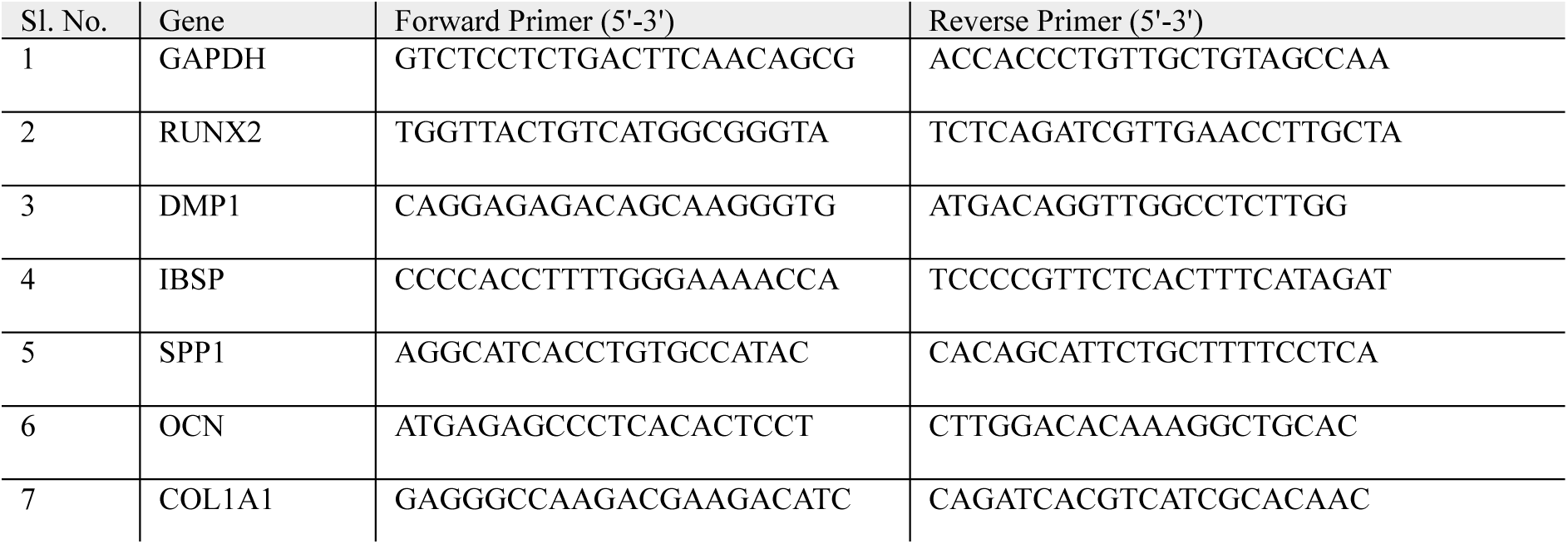
Primer sequences used for quantitative real-time PCR (qRT-PCR) analysis of osteogenic related genes.

## 6. Supporting information

Supporting Information is available from the Wiley Online Library or from the author.

## 7. Acknowledgment

This research was supported by the National Science Foundation through the NSF RUI grant (Award No. 2332041) and the Centre for Engineering Mechanobiology (CEMB, CMMI Award No. 15-48571), as well as by the National Institutes of Health through the University of Pennsylvania Institutional Research and Academic Career Development Award (IRACDA, K12GM081259), the Penn Center for Musculoskeletal Disorders (P30-AR069619), and the NIH R01 grant (1R01HD113596) from the National Institute of Arthritis and Musculoskeletal and Skin Diseases. The authors thank Dr. Robert Grabski (Research Scientist, University of Alabama at Birmingham) for his valuable assistance with confocal fluorescence microscopy imaging. The authors thank Dr. Emmanuel Tadjuidje (Professor Biology Department, Alabama State University) for the stereomicroscopic images. V.V. also gratefully acknowledges nTopology for providing an educational software license to Alabama State University.

## 9. Data availability statement

The data that support the findings of this study are available from the corresponding author upon reasonable request.

