## Supporting Information for "Plasma-Enabled Multiscale Coupling of Architecture and Biointerfaces Drives Osteogenesis in 3D-Printed Gyroid Scaffolds"

**Plasma-Enabled Multiscale Coupling of Architecture-Driven Mechanical Cues and Nanoscale Biointerface Engineering in Triply Periodic Minimal Lattice Gyroid Bone Scaffolds**

Pratheesh Kanakarajan Vijaya Kumari[1], Jasmine Carpenter[1], Barnett Cleon[2], Christopher J Panebianco[3], Joel D Boerckel[3], Derrick Dean[1], Vineeth M Vijayan[1]*

[1]Laboratory for Polymeric Biomaterials, Department of Biomedical and Mechanical Engineering, Alabama State University, Montgomery, Alabama 36104, United States of America

[2]Department Forensic and Physical Science, Alabama State University, Montgomery, Alabama 36104, United States of America

3Department of Bioengineering and Department of Orthopedic Surgery, University of Pennsylvania, Philadelphia, Pennsylvania 19104, United States of America

*This manuscript is currently under review in Advanced Healthcare Materials.*

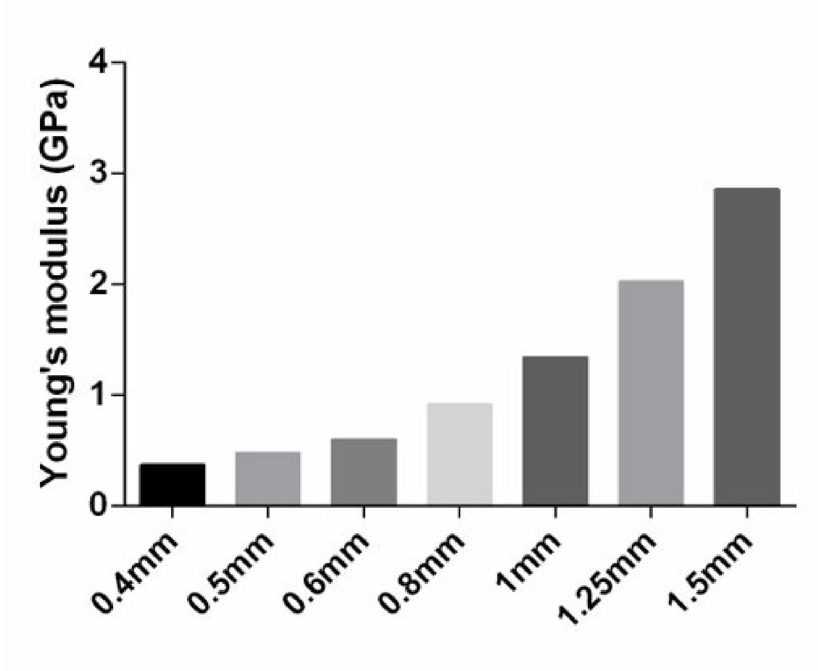


**Figure S1:** FEA predicted Young's modulus of TPMS gyroid scaffolds as a function of strut thickness, showing a progressive increase in stiffness with increasing strut size.

**Table S2.** Quantitative finite element analysis (FEA)–derived von Mises stress metrics across gyroid scaffold designs with varying strut thickness. Mean, standard deviation (SD), and maximum stress values were calculated from spatially sampled regions within each scaffold. The ratio of maximum-to-mean stress provides an indicator of stress concentration and heterogeneity within the architecture.

| Lattice Thickness (mm) | Mean Stress (Pa) | Standard Deviation (SD) (Pa) | Max Stress (Pa) | Max / Mean Stress |
|---|---|---|---|---|
| 0.4 | $9.69 \times 10^4$ | $5.19 \times 10^4$ | $1.49 \times 10^5$ | 1.54 |
| 0.6 | $7.61 \times 10^4$ | $4.28 \times 10^4$ | $1.23 \times 10^5$ | 1.62 |
| 0.8 | $4.95 \times 10^4$ | $3.10 \times 10^4$ | $8.99 \times 10^4$ | 1.82 |
| 1.0 | $3.69 \times 10^4$ | $2.85 \times 10^4$ | $8.21 \times 10^4$ | 2.22 |
| 1.25 | $2.15 \times 10^4$ | $1.74 \times 10^4$ | $4.37 \times 10^4$ | 2.03 |
| 1.5 | $1.60 \times 10^4$ | $1.45 \times 10^4$ | $4.21 \times 10^4$ | 2.63 |

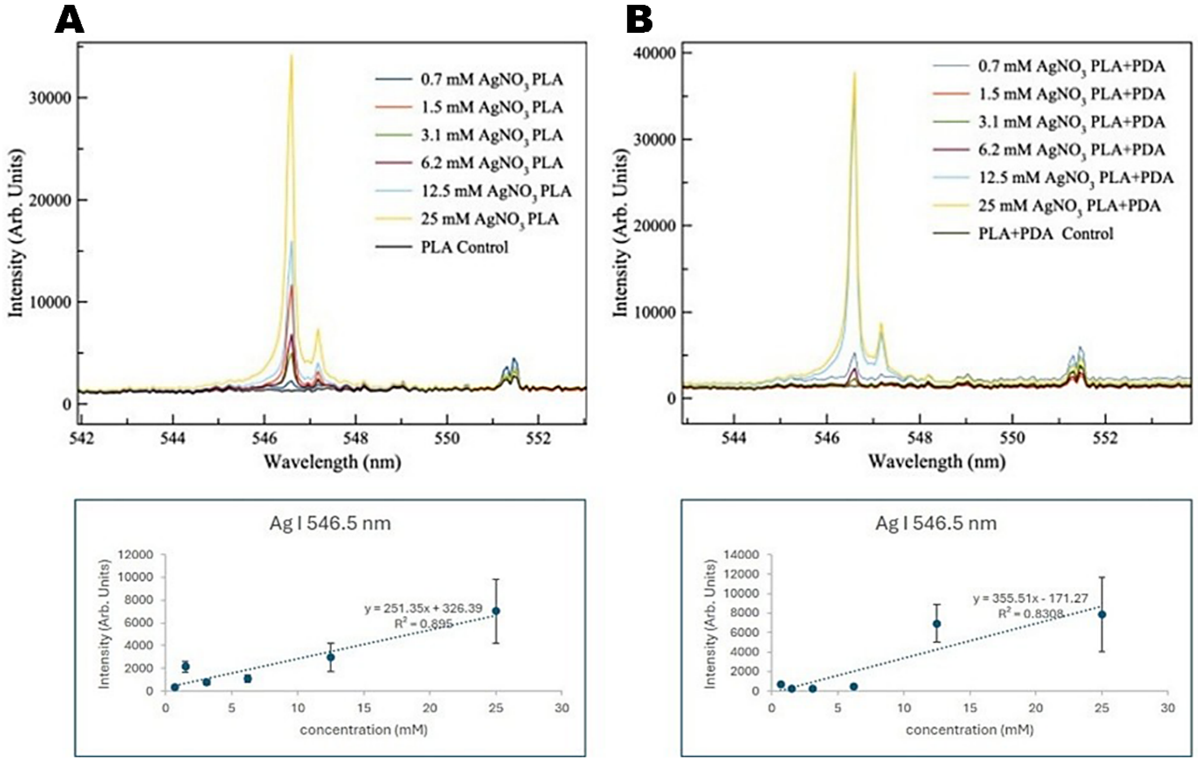


**Figure S3:** Laser-induced breakdown spectroscopy (LIBS) analysis of silver deposition on PLA and PDA-coated PLA scaffolds treated with varying $AgNO_3$ concentrations. Representative LIBS spectra showing the Ag I emission peak at 546.5

nm for PLA and PLA+PDA scaffolds, along with corresponding calibration plots correlating Ag I peak intensity with precursor concentration.

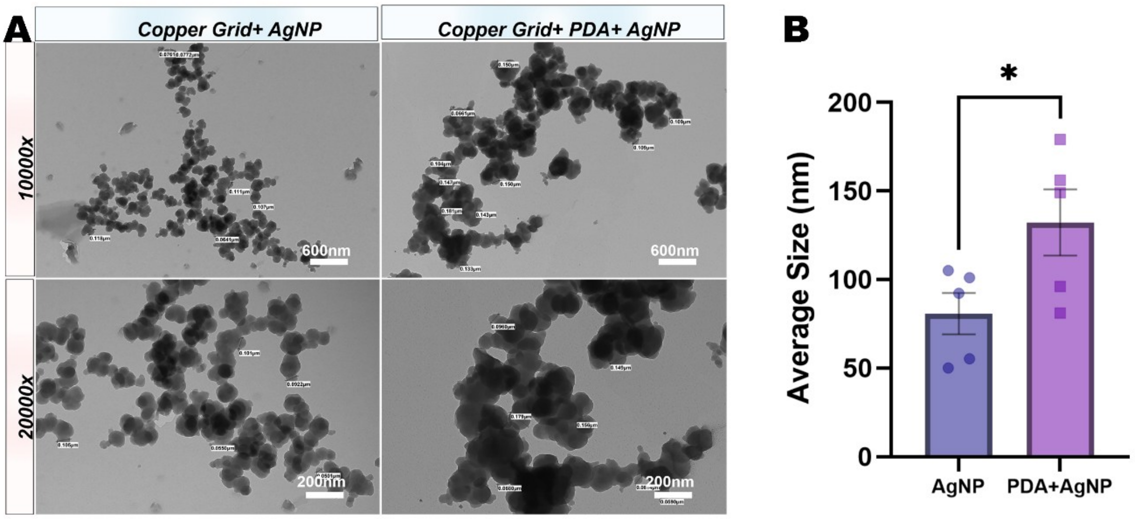


**Figure S4:** Transmission Electron Microscopy (TEM) Images of Silver Nanoparticles (AgNPs) on Copper Grids with and without Polydopamine (PDA) Coating.

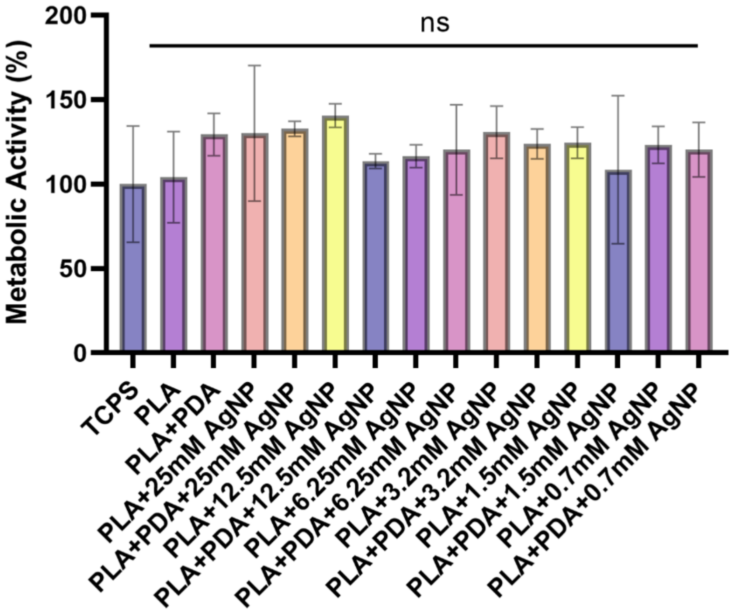

**Figure S5:** Alamar Blue assay showing metabolic activity of cells cultured on PLA, PDA coated, and AgNP modified scaffolds after 24 h. Data are presented as mean ± SEM, with no significant differences observed among groups (ns), indicating cytocompatibility across all conditions.

.

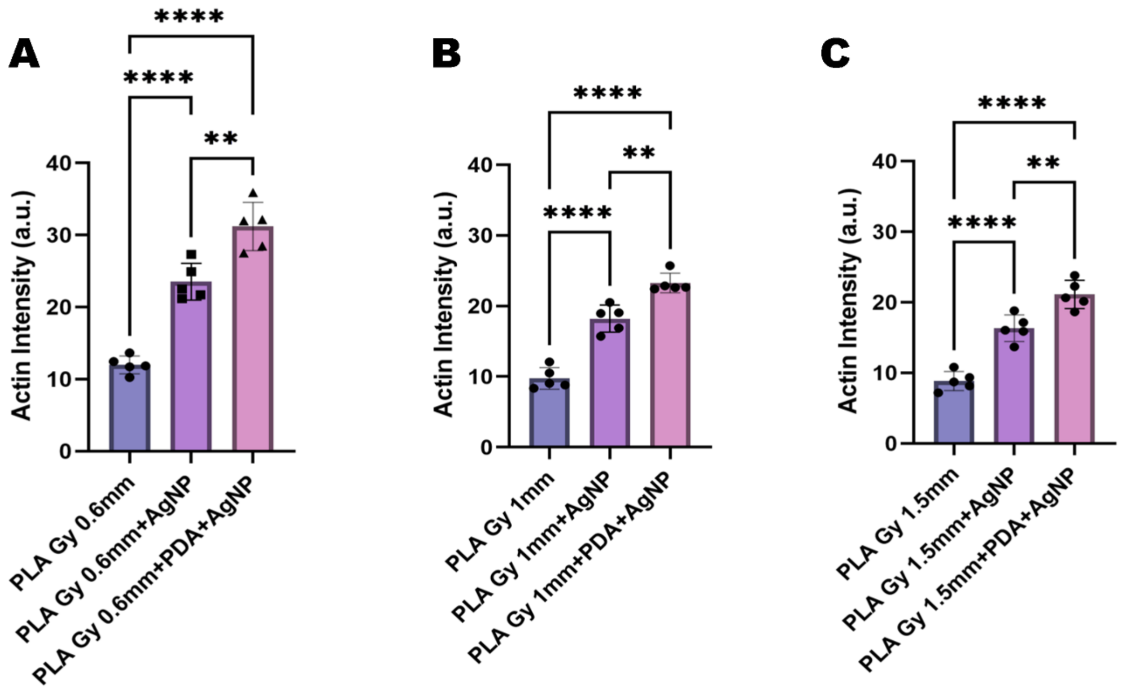


**Figure S6**: Quantitative analysis of actin fluorescence intensity in cells cultured on AgNP-functionalized PLA gyroid scaffolds. Actin intensity (a.u.) measured in cells seeded on PLA Gy, PLA Gy+AgNP, and PLA Gy+PDA+AgNP scaffolds at unit cell sizes of 0.6 mm, 1 mm, and 1.5 mm. Data are presented as mean ± SEM ($n = 5$). Statistical comparisons were performed using one-way ANOVA with Tukey's multiple comparisons test (** $p < 0.01$, **** $p < 0.0001$).
